# A structure-guided approach to non-coding variant evaluation for transcription factor binding using AlphaFold 3

**DOI:** 10.1101/2025.03.20.644433

**Authors:** Lukas Gerasimavicius, Simon C. Biddie, Joseph A. Marsh

## Abstract

Non-coding single-nucleotide variants (SNVs) that alter transcription factor (TF) binding can affect gene expression and contribute to disease. Sequence-based methods can excel at predicting TF binding, but rely on training data and can exhibit TF-specific biases. Here we propose a structure-guided approach for non-coding SNVs, using AlphaFold 3 (AF3) to model TF-DNA complexes and FoldX for downstream physics-based assessment. Benchmarked against SNP-SELEX data for six TFs (SPIB, ELK3, ETV4, SF-1, PAX5 and MEIS2), the FoldX-based strategy showed good agreement with experimental allele preferences. Interestingly, differences in AF3’s interface predicted template modelling (ipTM) score aligned even more closely with SNP-SELEX results, generally surpassing energy-based metrics. Application to known disease-associated variants recapitulated most reported effects for TFs including NKX2-5, GATA3 and USF2A-USF1. In these examples, considering both ΔipTM and FoldX energies proved more reliable than either metric alone. While less accurate than state-of-the-art sequence-based methods, this work demonstrates that structural modelling can yield interpretable insights into how non-coding variants influence TF binding. By highlighting both the promise and limitations of AF3 in this context, our study provides a framework for complementary structural evaluation of regulatory variants.

## Introduction

Transcription factors (TFs) are regulatory proteins that recognize and bind short DNA sequence motifs in *cis*-regulatory elements, modulating downstream gene expression, and thus controlling diverse cellular processes including development and responses to environmental cues (1). As such, single nucleotide variants (SNVs) affecting TF-DNA binding have been implicated in human disease (2–5). Missense variants can affect TF function directly by either entirely destabilizing the protein or more subtly affecting its binding preference(6). For example, missense variants in PAX6, an essential TF in eye development, can affect its DNA-binding specificity and affinity, leading to severe eye malformations (7, 8). However, expression of genes is also affected by mutations in the non-coding genome, which can disrupt existing enhancer and promoter motifs, or enable new TF binding sites by affecting chromatin accessibility or generating functional binding motifs (9–11).

While most disease variants in databases like ClinVar (12) are primarily associated with simple Mendelian diseases, non-coding variation is significantly more difficult to causally associate and characterize. Most SNVs linked to common disease and complex phenotypes through genome-wide association studies (GWAS) occur in non-coding regions, with many variants contributing small effect sizes (13, 14). Over 90% of such GWAS SNVs fall into non-coding regions, with ∼70% estimated to overlap TF binding motifs (15, 16). However, clinical examples of regulatory variants directly leading to detectable disease through interplay with TFs are scarce. Despite efforts, the direct functional consequences of most disease-associated non-coding SNVs from GWAS remain unelucidated.

To tackle this issue, numerous high-throughput experimental approaches have been developed to examine TF-DNA binding and the effects of SNVs on these interactions. *In vivo* experiments such as chromatin immunoprecipitation sequencing (ChIP-seq) can identify TF binding sites, in context with epigenetic and cell type-specific factors like chromatin accessibility, which govern available TF binding sites (17). ChIP-seq also enables the comparison of TF allelic binding preference, facilitating the evaluation of SNV effects (18). However, TF binding signals identified by ChIP-seq can span hundreds of nucleobases (16), while TFs are known to bind more specific 6-20 base motifs (19). Binding sites can be identified at a higher resolution through *in vitro* methods such as protein binding microarrays (20) or systematic evolution of ligands by exponential enrichment (SELEX) experiments (21, 22). An iteration on the latter, termed SNP-SELEX, has recently been utilized to compare TF-specific binding events between reference and SNV DNA sequences, constrained to a 40 base resolution (23). All of these approaches have contributed to the identification of TF binding motifs in database resources such as JASPAR (24), ENCODE (25), ADASTRA (18) and ANANASTRA (26).

Despite the success of these methods, experimental approaches are often time-consuming and expensive to perform at scale, which has led to a massive effort in developing computational methodologies to predict sequences bound by TFs and the effects of non-coding SNVs. Position weight matrices (PWMs) are among the simplest methods for evaluating TF binding (27). Utilising experimental data to generate TF-specific motif binding logos, PWMs represent the nucleotide probabilities at each position within a sequence likely to be bound by a given TF. However, far more advanced strategies have also been developed, such as gapped k-mer approaches like deltaSVM (28), which can capture nucleotide dependencies and have been recognised as top-performing methods in benchmarks (29). Deep learning has also been applied to TF binding site prediction, with methods like DeepSEA (30) being shown to more accurately predict *in vivo* binding by incorporating complex features like chromatin accessibility (16). Although these approaches differ in their training data and the scope of their feature engineering, they generally all rely on sequence information.

While sequence-based approaches have achieved significant success (29, 31), they cannot assess the structural complexity of TF-DNA interactions, which could provide deeper mechanistic insights into binding specificity and the effects of genetic variants. Previous studies have demonstrated that the effects of missense mutations in TFs can be accurately assessed using structure-based stability predictors, particularly when a TF-DNA complex structure is available (32). FoldX, a computational stability prediction tool, evaluates effects of missense mutations, but, additionally, has the functionality to mutate DNA and RNA sequences in a complex structure, allowing assessment of non-coding SNVs effects on TF binding (33–35). However, the major hurdle up until recently has been a lack of viable TF-DNA structures in existing databases, and limitations of structure modelling tools. With the recent release of AlphaFold 3 (36) (AF3) and similar methods (37, 38), we are now able to predict biomolecular complex structures, integrating proteins, DNA, RNA and ligands within a single model. This paves the way for using downstream structure-based methods to assess the accuracy of TF-DNA binding predictions against extensive TF binding datasets.

In this work we adapt our previous structure-based methodology, used for missense variant evaluation, to assess the impact of non-coding SNVs on TF binding. We leverage AF3 to derive TF-DNA complex models and use SNP-SELEX data as ground truth for assessing the accuracy of structural TF binding preference predictions. This dataset provides a simplified binding scenario, free from confounding factors such as DNA methylation and chromatin accessibility, making it well-suited to directly evaluate the utility of stability predictors like FoldX. We derive a selection of structure- and physics-based metrics. We show that FoldX predictions can recapitulate SNP-SELEX assay results to a high degree. However, we find that the AF3 interface predicted template modelling (ipTM) score, a byproduct of deep-learning, often performs better in identifying differentially bound sequences. We demonstrate high performance heterogeneity across the tested TFs, and that FoldX predictions can outperform ΔipTM in specific cases, providing orthogonal support. This heterogeneity suggests TF-specific modelling strategies may be required, such as focusing on binding domains or considering cooperative TF assemblies.

## Results

### Structural metric distributions recapitulate qualitative SNP-SELEX binding groups

We set out to test whether structural modelling can capture differences in transcription factor (TF) binding induced by single-nucleotide variants. SNP-SELEX provides an ideal benchmark, as it reports preferential binding scores (PBS) for paired reference (*ref*.) and alternative (*alt*.) sequences *in vitro*. To keep the analysis tractable while covering diverse scenarios, we selected six TFs: three ETS family members with well-defined monomeric binding (SPIB, ETV4, ELK3) (39), SF-1 from the nuclear receptor family (reported to bind DNA as a monomer) (40), PAX5 (modelled with its DNA-binding domain only, due to poor performance of full-length assays), and MEIS2, a homeobox protein known to act cooperatively (41, 42). This panel allowed us to test the approach across both straightforward and more challenging binding contexts.

For each reference-alternative allele pairs, we generated AF3 models of the TF-DNA complex, covering both reference and alternative alleles, totalling 75,134 modelled complexes. We then used FoldX to assess the impact of SNVs on the overall stability of the complexes, and derived two structural impact scores. The first approach involved calculating a ‘symmetric’ ΔΔG value (ΔΔG_sym_) as the mean of both the forward and reverse DNA mutations. The ΔΔG from the reference allele structure will not necessarily be the inverse of the ΔΔG calculated from the alternative allele structure, considering there may be a binding site difference between the two predicted models. The second approach, ΔInterface, directly evaluates the difference in interaction energy between the protein and the DNA chains for the two structures. Finally, we were also curious whether AF3, as a result of its extensive training on evolutionary information, biomolecular structures, and specific examples of TF-bound DNA sequences, will have learned a sequence preference representation. To assess this, we used the AF3 interface predicted template modelling (ipTM) score, a confidence measure for the relative positions of complex subunits to derive ΔipTM, as the difference between reference and alternative model confidences.

To evaluate the performance of our metrics, we used the allele preference classification outlined in the SNP-SELEX study. Each oligomer pair, differing by a single central nucleotide, was designated as either non-preferentially bound (non-pbSNP) if no significant differential binding was observed, or preferentially bound (pbSNP), where one allele was significantly favoured. pbSNPs were further classified by direction of preference, towards either the *ref*. or *alt*. allele. Ideally, non-pbSNP pairs should yield metric values close to zero, while pbSNPs should show large positive or negative values, indicating *ref*. or *alt*. preferences, respectively.

Pooling data across TFs, **Figure 1a** demonstrates that all three metrics separated the SNP-SELEX preference groups, with highly significant differences between the *ref*., *alt*. and non-pbSNP groups. Score directions also matched expectations. However, the methods do show performance variability, with ΔΔG_sym_ demonstrating an overall shift towards more positive Δ values for all three allele preference groups, corresponding to more stable interactions in the reference allele models. ΔInterface produced a more balanced distribution, centered near zero for non-pbSNPs. This difference likely arises because ΔInterface captures only direct protein-DNA interaction energies, whereas ΔΔG_sym_ can be influenced by AF3 placing the TF way from the variant site, diminishing the contribution from one allele model.

**Figure 1.**
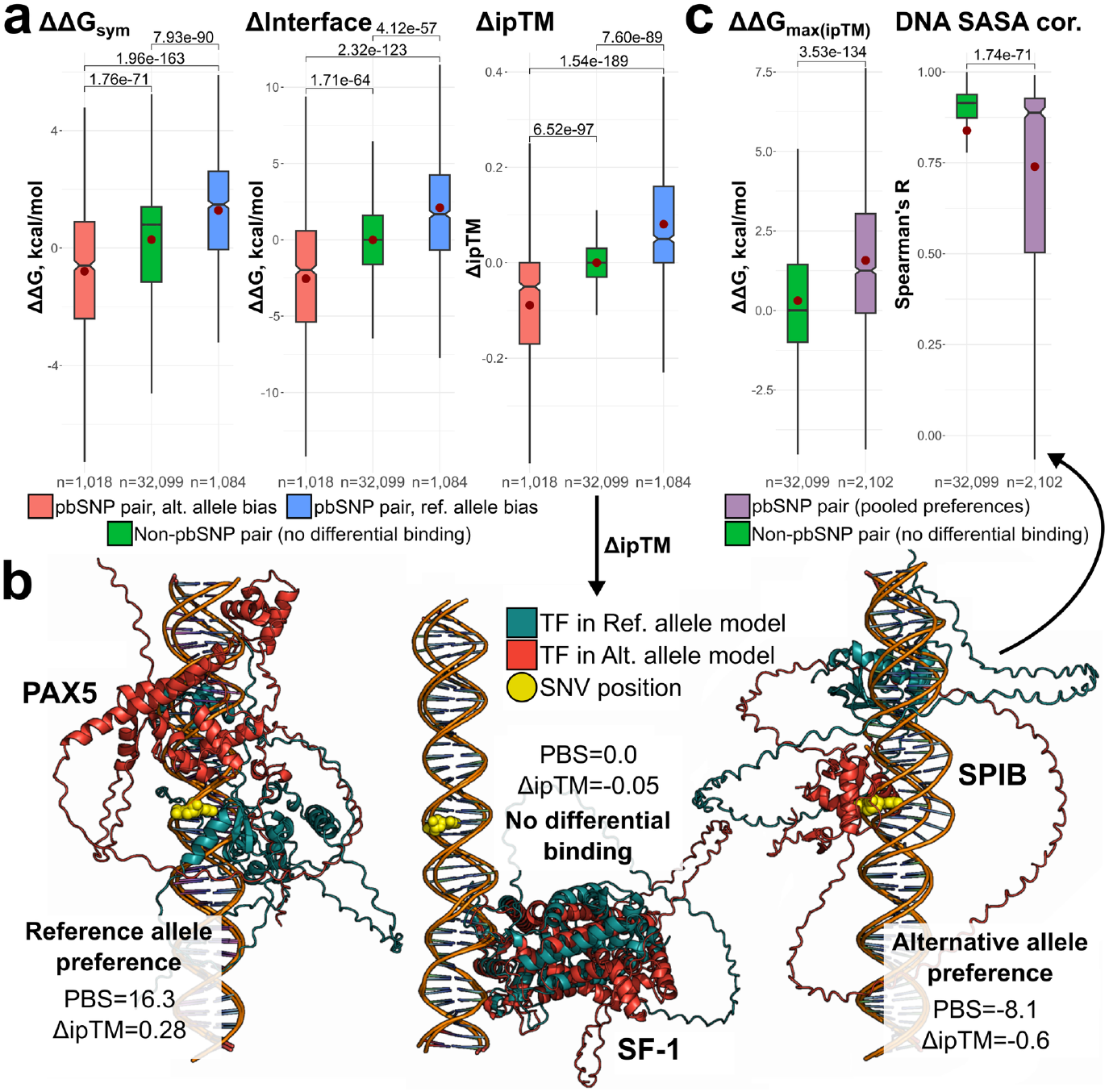
Structure-derived TF-DNA stability metrics qualitatively recapitulate SNP-SELEX differential binding results. Analysis represents allele pairs pooled from all six TFs: SPIB, ETV4, ELK3, SF-1, PAX5 and MEIS2. **a** – Structural metric score distributions across SNP-SELEX allele preference groups. **b** – AlphaFold 3 can provide highly interpretable predictions for assessing the impact of non-coding SNVs on transcription factor binding. Cyan TF chains represent the binding pose and site for the reference DNA sequence, while the red structures show the predicted binding site for the alternative allele. The yellow orbs represent SNV location on the forward strand. PBS – SNP-SELEX preferential binding scores. ΔipTM – difference in the AF3 interface predicted template modelling score between the reference and alternative allele models. Positive values indicate reference allele preference, negative values indicate binding bias for the alternative allele, while values close to zero indicate no differential binding. **c** – Structural metrics inspired by AF3’s binding site shifts show discriminatory power between pbSNP and non-pbSNP pairs. ΔΔG_max(ipTM)_ metric was derived for the model with higher ipTM of the pair. DNA SASA cor. represents Spearman’s correlation between per-nucleotide DNA solvent accessible surface areas (SASA) between the paired models. (**a**,**c**) Boxes denote data within 25th and 75th percentiles, and contain median (middle line) and mean (red dot) value notations. Whiskers extend from the box to furthest values within 1.5x the inter-quartile range. Significance values are derived using two-sided Holm-corrected Dunn’s tests (**a**) or two-sided Mann-Whitney *U* tests (**c**). Sample size (n) indicates the number of allele pairs.

Surprisingly, ΔipTM gave the best overall performance, with the strongest separation of preference groups, and the tightest distribution for non-pbSNPs. AF3 byproduct metrics have been previously shown to capture proxy information on the impacts of missense variants on biophysical stability(43). Here, the relative difference in ipTMs also reflects non-coding variant binding preference. Nonetheless, while the two FoldX scores lead to poorer group separation, they are both driven and constrained by the underlying AF3 binding site prediction, and provide additional noise through structure relaxation and sidechain adjustment. From this perspective, the ability of the FoldX scores to maintain the overall correct directionality of the predicted TF preference, with statistically significant distinction for qualitative preference groups, is reassuring both in terms of FoldX applicability and TF placement by AF3.

Notably, all the metrics display TF-specific performance heterogeneity. **Supplementary Figure 1** reveals that ΔipTM performs well for ETS TFs like SPIB, as well as SF-1, in discriminating between the qualitative binding groups. Score overlap increases for PAX5, but differences remain significant. However, ΔipTM fails to differentiate between the allele preference groups for MEIS2. This could potentially be influenced by most DNA-binding MEIS structural templates from the Protein Data Bank (PDB) being dimeric or cooperative (42, 44), or the SNP-SELEX experiment representing cooperative binding results, whereas we modelled monomeric TF binding. Interestingly, FoldX-derived scores showed a somewhat different pattern. While ΔInterface predictions corresponded with ΔipTM (**Supplementary Figure 2, a**), ΔΔG_sym_ values separated MEIS2 preference groups but performed poorly on PAX5 (**Supplementary Figure 2, b**). As AF3 depends on training examples, both for overall protein structure prediction and DNA sequence interactions, models for some TFs may not be as accurate due to fewer available TF-DNA complexes in the PDB, or lower quality binding motif data in JASPAR, which was used to positively reinforce the model through generated DNA sequence examples. The observed performance heterogeneity suggests that considerations should be taken when utilizing AF3 to predict TF-DNA complex models, such as ensuring the functionally relevant multimeric state is being modelled, and utilizing multiple metrics. Although ΔipTM generally gave the strongest group separation, FoldX metrics may retain value in specific cases.

### AlphaFold 3 tends to predict shifted TF binding sites for differentially bound pairs

Given the performance of ΔipTM, and the structural nature of the approach, we next examined AF3 models for selected SNV pairs where metric scores matched the SNP-SELEX allele preference groups. **Figure 1b** shows examples from each class: a variant with diminished TF binding (*ref*. allele preference), a neutral variant (non-pbSNP), and a variant enhancing TF binding (*alt*. allele preference). For the non-pbSNP case, AF3 positioned the TF away from the variant in both models, consistent with the lack of binding difference observed. In preferential cases, at least one modelplaced the TF in contact with the variant, shifted way in the *ref*. allele preference, or shifted towards the SNV where the *alt*. allele provided a higher confidence binding motif.

While AF3 often predicts distinct binding shifts for pbSNP pairs, these variants are more likely to alter the affinity of the same binding site rather than create new ones. Our results are based on single-seed AF3 predictions, which yield only the most probable TF position for a given input, and do not capture the broader landscape. The true binding sites on the 40 bp SNP-SELEX oligomers are unknown and cannot be resolved with certainty without further experiments. Thus, AF3 predictions should not be taken as literal representations of TF binding modes. Instead, our goal is to assess their behaviour and identify practical use cases for AF3 complex modelling.

Because only a single nucleotide is substituted, the large binding site shifts predicted by AF3 are surprising. We speculated this may partly underlie the performance of the FoldX metrics. To test this, we explored additional structure-derived metrics. First, we used the FoldX ΔΔG from the model structure with the higher ipTM in each pair (ΔΔG_max(ipTM)_), combining AF3 confidence in binding placement with direct energetic evaluation. This metric distinguished between the two pair groups effectively, as the higher ipTM models of pbSNPs tended to place the TF consistently at the variant site, yielding a non-zero ΔΔG value (**Figure 1c**). Second, we asked whether metric performance simply reflected AF3 predicting distinct TF binding sites between alleles. To assess this, we compared nucleotide solvent accessible surface area (SASA) between allele models using Spearman’s correlation (*DNA SASA cor*.). High correlation indicates similar TF positioning, while low correlation suggests SNV-induced shifts. As expected, pbSNPs showed variable correlations, while non-pbSNPs remained highly correlated (**Figure 1c**).

We next examined whether AF3-predicted binding site shifts fully explained metric performance. Histograms of *DNA SASA cor*. values showed a bi-modal distribution for both pbSNPs and non-pbSNPs, with most allele pairs highly correlated (>0.8), but a larger fraction of low-correlation cases among pbSNPs (**Supplementary Figure 3a**). Separating the data at this 0.8 threshold, we found that structural metrics still recapitulated SNP-SELEX classifications, even when TF positions were aligned, indicating that site shifts are not the sole performance driver (**Supplementary Figure 3b**). While some metrics showed stronger separation in the low-correlation subset, ΔΔG_sym_ displayed very similar group means (-0.81 *vs* -0.74 for *alt*. preference, 0.28 *vs* 0.33 for non-pbSNPs and 1.29 *vs* 1.28 for *ref*. preference), suggesting little dependence on positional changes. TF-specific effects were also evident: PAX5 showed no significant difference in shift distributions between pbSNPs and non-pBSNPs, whereas MEIS2 exhibited a marked distribution shift yet also showed the weakest metric performance (**Supplementary Figure 4**). These results indicate that favourable use of structural metrics cannot be attributed solely to AF3 predicting shifted binding sites, but, rather, reflects AF3’s broader capacity to capture aspects of TF-DNA recognition and variant impact.

### Structural metrics are better at identifying preference direction than differential binding

Having established that structural metrics broadly recapitulate qualitative SNP-SELEX preference groups, we next evaluated their quantitative performance. Using the full dataset, including unclassified intermediate allele pairs, we assessed the overall correlation between PBS values and our three metrics (**Supplementary Figure 5**). Despite the agreement with qualitative classifications, Spearman’s correlations across the full dataset were weak, and even insignificant for PAX5 and MEIS2.

To explore perfomance in more detail, we first assessed how well structure-derived metrics distinguish pbSNPs from non-pbSNPs. For this analysis, we used absolute values of ΔΔG_sym_, ΔInterface and ΔipTM, as well as raw ΔΔG_max(ipTM)_ and *DNA SASA cor*., since these latter metrics do not distinguish between *ref*. and *alt*. preference by default. **Figure 2a** demonstrates receiver operating characteristic (ROC) curves for data pooled from the six TFs, with the area under the curve (AUC) indicating the performance of each metric. ΔipTM performs best (AUC = 0.685), followed by ΔΔG_max(ipTM)_, which outperformed its symmetric counterpart. The ROC curve for ΔΔG_sym_ also shows that many pbSNPs were assigned low differential binding scores. Interestingly, precision-recall (PR) curves (**Supplementary Figure 6a**), which weigh the accurate identification of pbSNPs more than non-pbSNPs, indicate that both of the ΔΔG metrics perform better.

**Figure 2.**
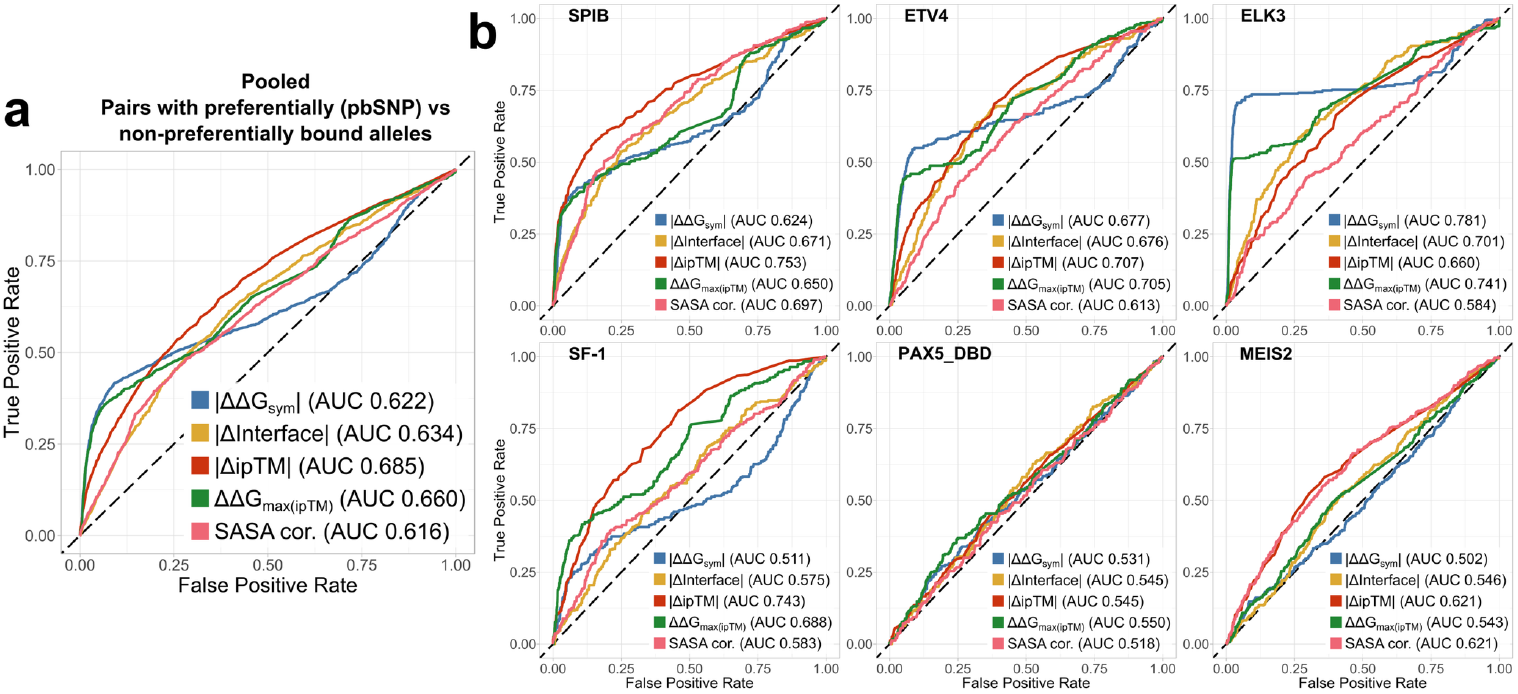
Structure-derived allele preference metrics show capacity to distinguish between differentially bound allele pairs, but the performance is highly heterogenous across TFs. Analyses represent receiver operating characteristic (ROC) curves, with area under the curve (AUC) representing the classification performance. **a** – Generalized metric performance in distinguishing between pbSNP containing pairs and non-differentially bound sequences (non-pbSNPs). Analysis represents allele pairs pooled from all six TFs: SPIB, ETV4, ELK3, SF-1, PAX5 and MEIS2. **b** – TF-wise classification performance. Qualitative binding preference groups were annotated in line with the SNP-SELEX paper classification. Structural metric directions were adjusted to match the directional convention in the analyses.

Exploring performance by TF revealed substantial heterogeneity (**Figure 2b**). The highest overall AUC was achieved by ΔΔG_sym_ on ELK3 (0.781), outperforming ΔipTM for the same TF (0.660). By contrast, ΔipTM reached its best performance on SPIB (0.753), followed by *DNA SASA cor*. (0.697), suggesting that AF3 predicts shifted binding sites more readily for some TFs than others. For PAX5, none of the metrics performed well, with the highest AUC only 0.550 for ΔΔG_max(ipTM)_. ΔipTM and *DNA SASA cor*. showed moderate performance for MEIS2 (0.621). Similar trends are mirrored by the PR analysis (**Supplementary Figure 6b**). Overall, these results suggest that the best structural metric is TF-dependent, though pooled analyses suggest ΔipTM is the most generalizable.

It is also crucial to assess how well the metrics identify the correct preference direction in differentially bound pbSNP pairs. **Figure 3a** shows that all three metrics are performing much better at this task, with ΔipTM demonstrating an AUC of 0.792, only slightly surpassing ΔΔG_sym_, and is also mirrored by PR analysis (**Supplementary Figure 7**). For most TFs, ΔipTM is the top predictor, with ELK3 and MEIS2 being the only outliers where ΔΔG_sym_ provides more accurate predictions (**Figure 3b**). However, PR analyses do indicate prediction performance bias for specific directions and TFs, with *ref*. (MEIS2, PAX5, SF-1) or *alt*. (SPIB) preference being more often accurately predicted (**Supplementary Figure 8**). The robustness of ΔipTM is also corroborated by Matthews correlation coefficient, which assesses the accuracy of prediction direction (**Figure 3c**).

**Figure 3.**
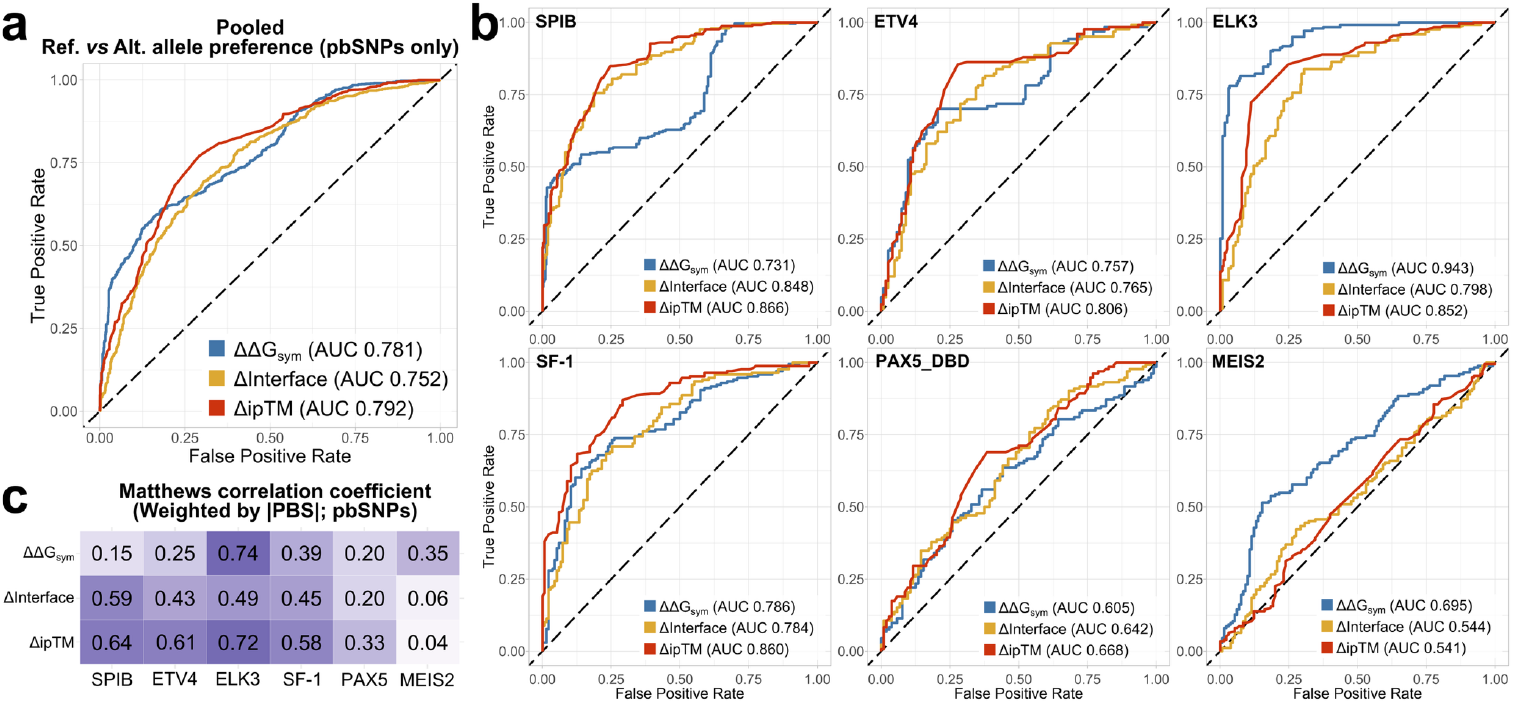
Structural TF binding metrics are considerably better at identifying the preferred allele in differentially bound allele pairs than distinguishing the fact of preference itself. Analyses represent receiver operating characteristic (ROC) curves, with area under the curve (AUC) representing the classification performance. **a** – Generalized metric performance in identifying the preferred allele from among differentially bound sequences (pbSNP pairs only). **b** – TF-wise classification performance. Qualitative binding preference groups were annotated in line with the SNP-SELEX paper classification. Structural metric directions were adjusted to match the directional convention in the analysis. **c** – TF-wise Matthews correlation coefficients for structural metric accuracy. Each cell reflects agreement between the metric’s sign and the experimental PBS sign, with variants weighted by their |PBS| (stronger experimental effects have greater influence; near-zero PBS contribute minimally).

While the structural metrics show some utility in identifying preferentially bound alleles, high-performance sequence-based methodologies already exist. DeltaSVM (28) represents state-of-the-art prediction in TF binding preference, and three of our tested TFs have validated high-quality deltasVM models, trained on HT-SELEX data (18), although we do note that the SNP-SELEX data we use here was also used for deltaSVM model selection (23). **Supplementary Figure 9** demonstrates that deltaSVM demonstrates nearly perfect prediction across ELK3, SPIB and SF-1 at both classification tasks. At the same time, out of the 533 unique transcription factors for which HT-SELEX data was used to derive deltaSVM models, only 94 made it into the high-confidence category, underscoring both the dependence on experimental assays and their variability. By contrast, our structural framework requires no additional training data and could, in principle, be applied broadly, including to TFs and variant contexts beyond the reach of existing sequence-based predictors.

### ipTM magnitude is not an indicator of ΔipTM performance

While ΔipTM can predict DNA sequence preferences, its practical use depends on knowing which complex predictions are reliable without experimental validation. In our benchmarking, we used SNP-SELEX sequences, all confirmed as containing TF binding sites, yet many corresponding AF3 models fell below the ipTM confidence threshold of 0.6 (**Figure 4a**), which is generally indicative of a low confidence model and a likely failed prediction (45, 46). Moreover, ipTM magnitude did not correlate with metric performance: AUROC values for pbSNP versus non-pbSNP classification showed no significant association (**Figure 4b**). For example, SF-1, one of the best-performing TFs by structural metrics, had an average ipTM of only 0.51, whereas MEIS2 had the third-highest average ipTM but poor metric performance.

**Figure 4.**
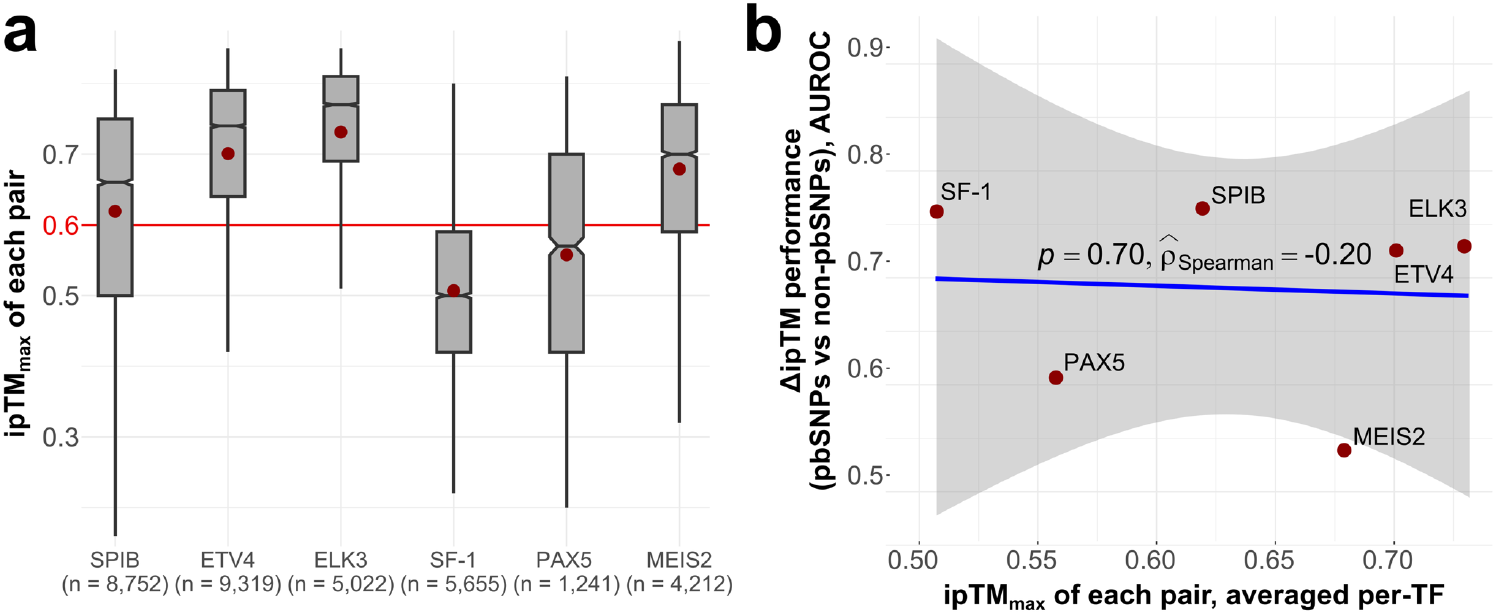
ipTM magnitudes are not indicative of the underlying model utility and ΔipTM performance. **a** – distributions of model ipTMs across TFs. The maximal ipTM of each pairs is reported. Boxes denote data within 25th and 75th percentiles, and contain median (middle line) and mean (red dot) value notations. Whiskers extend from the box to furthest values within 1.5x the inter-quartile range. Sample size (n) indicates the number of allele pairs. **b** – Correlation between mean ipTM per-TF and the ΔipTM performance in distinguishing pbSNPs from non-pbSNPs. Rho represents Spearman’s correlation.

Issues have been noted with the way the absolute ipTM values are calculated and how they can be artificially inflated, and alternative quality metrics have been devised (47, 48). The metrics improved correspondence with experimental results for prioritizing peptide binding, so we tested whether a similar approach could help for TF-DNA complex models. We derived ipSAE (47) values, using either the mean of the largest inter-chain ipSAEs or the largest protein–DNA ipSAE alone. However, **Supplementary Figure 10** reveals that neither approach improved correspondence with experimental validation, and TF-specific patterns remained similar to ipTM. Other metrics may provide a better solution, or novel confidence scores tailored to models containing nucleic acids may need to be developed. Overall, AF3 and similar methodologies are known to underperform on nucleic acid complex, largely due to limited training examples in the PDB (49). Thus, the applicability of our framework to unvalidated binding events, and the threshold at which an ipTM value indicates true model failure, remain open questions.

### Clinically relevant non-coding SNVs can be accurately identified through structural metrics

To better assess the practical applicability of our metrics, we extended the analysis to published disease-associated non-coding SNVs (**Supplementary Table 1**). As no magnitude thresholds for preferential binding have been established, we applied a qualitative consensus approach, asking whether ΔΔG_sym_, ΔInterface and ΔipTM agreed in direction and showed shifts comparable to those observed in SNP-SELEX distributions.

A rare non-coding chr11:31685945 G>T variant has been shown to affect an autoregulatory PAX6 enhancer, lowering the binding affinity of PAX6 and disrupting the feedback loop crucial for correct eye development (50). However, our structure-guided approach failed to conclusively reproduce the experimentally validated effect of the variant, with the FoldX metrics either not showing a strong enough signal or indicating the incorrect allele preference. ΔipTM weakly suggested the variant reduces binding preference, despite both models essentially having the same binding site (**Figure 5a**). This example reflects a pattern of less accurate allelic preference prediction for PAX family TFs through structure-derived metrics, concerning both clinical and *in vivo* assay data. A possible explanation is that the JASPAR motifs, which were used to optimize AF3 for DNA binding, might not actually be representative enough of all cases, as deltaSVM PAX5 motif models trained on HT-SELEX (23) data failed to break 0.75 AUC in the SNP-SELEX experiment. Another possibility is that AF3 might be biased by the PDB templates, as the resulting models are very similar to the PAX6 PDB structure containing a DNA-bound PAX6 paired domain (**Supplementary Figure 11**). Upon complex structure alignment, the T-rich nucleotide sequence from the PAX6 structure overlaps the model DNA, with a T nucleotide being positioned exactly at the variant location, and T also being the alternative allele in the models, which may have inflated the ipTM value. Finally, ΔipTM and ΔInterface are not solely influenced by the interactions of protein and DNA at the variant location, but by all interfaces made, which may introduce additional noise for PAX TFs and similar proteins which have multiple DNA-interacting domains.

**Figure 5.**
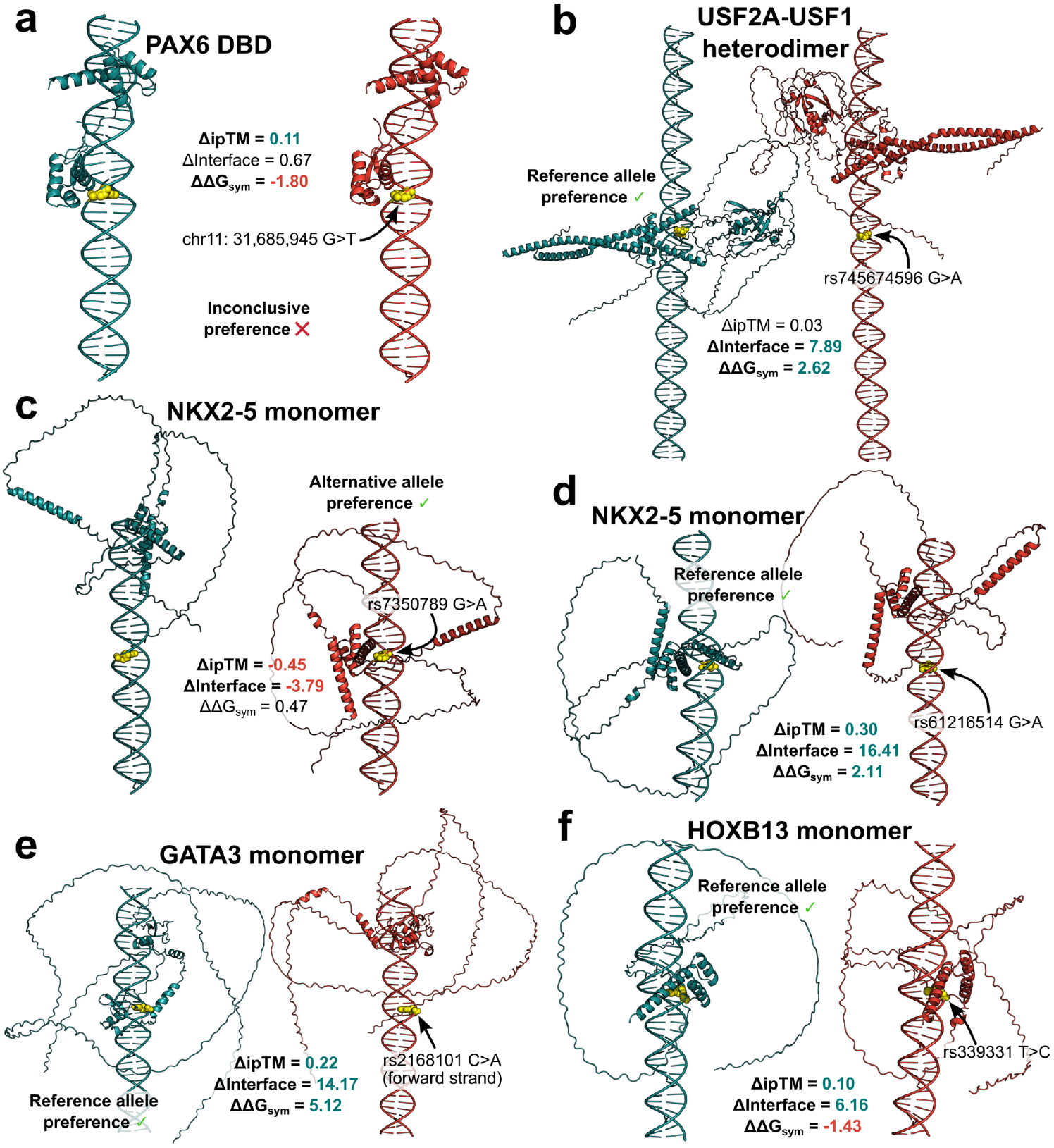
A consensus of structure-derived metrics recapitulates most tested clinical examples. Cyan models represent TF binding sites for the reference DNA sequence, while red structures show the predicted binding site for the alternative allele. Yellow orbs represent the SNV location on the forward strand. Structural complex models were predicted using the AF3 server. **a** – The structure-guided approach fails to conclusively identify differential binding direction for a regulatory PAX6 enhancer variant. **b** – FoldX protocols recapitulate assay data for a clinical variant affecting USF2A binding, where ΔipTM fails to inform of the effect. **c, d** – Structural metric consensus correctly identifies NKX2-5 allele preference for two SNV cases with opposite effects. **e** – Structure-guided metrics correctly predict the effects of an oncoprotective variant. **f** – A majority consensus approach of structural metrics can identify differential allele binding even without obvious binding site shifts.

Plaisancié *et al*. have recently characterized a regulatory rs745674596 G>A variant responsible for reducing the expression of the FOXE3 transcription factor, leading to ocular anomalies in a patient (51). The transcription factor USF2A was found to preferentially bind the reference allele, with reduced binding to the alternative allele sequence in a pull-down assay. We applied our approach to model the preferential binding of the USF2A-USF1 TF complex, as UniProt indicates that this is the predominant DNA-binding state. **Figure 5b** demonstrates that AF3 predicts distinct TF binding sites for the reference and SNV sequences. While the ΔipTM of 0.03 does not strongly suggest differential binding, the downstream FoldX predictions show a TF preference for the reference allele sequence. This example highlights how directional agreement across metrics can provide supportive evidence, even when single scores are weak. Notably, both predicted models had low overall confidence (ipTM 0.36 for the reference and 0.33 for the alternative).

We examined two NKX2-5 homeodomain variants, previously tested by electrophoretic mobility shift assays and implicated in congenital heart disorders, which altered binding in opposite directions (52). The predicted effects of rs7350789 G>A, which increased NKX2-5 binding, and rs61216514 G>A, which reduced it, are illustrated in **Figures 5c** and **5d**, respectively. AF3 predictions recapitulated these effects, with ΔipTM values matching the experimental results. Not relying on any single metric can help predict variant effects in line with the experimental data.

Non-coding variants in regulatory elements can drive cancer susceptibility, but some also act protectively by reducing oncogene expression through impaired TF binding. LMO1, a transcriptional co-factor implicated as an oncogene in neuroblastoma, provides one such example. The rs2168101 C>A variant in an LMO1 enhancer disrupts GATA3 binding, as shown by ChIP-seq experiments (53). Our structural metrics reproduced this effect, with all three agreeing on reduced binding of the SNV sequence (**Figure 5e**). We also examined rs339331 T>C, an oncoprotective variant in an RFX6 enhancer, where upregulation of RFX6 is associated with prostate cancer progression and metastasis (54). While AF3 predicted HOXB13 binding at the variant site in both alleles, two of the three metrics correctly indicated loss of preference for the alternative allele, consistent with its protective role (**Figure 5f**).

## Discussion

We set out to address the challenge of evaluating non-coding SNVs that influence TF binding and gene expression using structure-based approaches. With the advent of AF3, large numbers of TF-DNA complexes can now be modelled, making such an analysis feasible (36–38, 55). Although the structure-derived metrics we tested did not match the accuracy of a state-of-the-art sequence-based predictor (28), our results serve as a proof-of-concept that structure-based approaches can yield mechanistic insight and interpretable predictions of regulatory variant effects without specialized training.

The novelty of this framework is underscored by several observations. Notably, ΔipTM, as a byproduct of evolutionary, structural and DNA sequence motif training, emerges as the most consistent indicator of differential TF binding out of our derived metrics. AF3’s altered confidence in response to SNVs suggests that the model has learned a representation of DNA sequence preference. FoldX-derived scores often captured the correct direction of preference and outperformed ΔipTM for some TFs, such as ELK3 and MEIS2. The orthogonality of these two metrics implies that consensus evaluation can provide more robust predictions than either alone. Importantly, application to clinically reported variants demonstrated that structural metrics recapitulate experimentally validated effects in most cases, underscoring the translational potential of the approach.

Numerous sequence-based TF binding predictors exist, ranging from the simplest PWMs (16, 56), to multi-feature deep learning (57) and gapped k-mer models trained on large amounts of experimental data (28), such as deltaSVM, which achieve near-perfect predictive accuracy for well-studied TFs with extensive training data. However, structural modelling offers several advantages that sequence-only methods cannot provide. Firstly, it allows visualization of putative TF-DNA interactions, offering mechanistic explanations for how variants influence binding rather than just statistical prediction. It also enables downstream physics-based evaluation with stability predictors, which require no TF-specific training and could enable immediate applicability to TFs that lack high-quality experimental data. While AF3 binding site choices depend on previous training, the probable binding site landscape could potentially be identified directly through stability predictors. For example, one could generate structures for shorter motif-length subsequences within an oligomer and compare their energetics, thereby reducing reliance on AF3’s initial placement. Finally, structure-guided approaches can be applied beyond simple monomeric TF interactions, extending to homomeric and heteromeric TF complexes. While currently underpowered, a structural approach could complement existing sequence-based predictors by providing additional mechanistic interpretability, overall improving our ability to assess non-coding variant effects.

Nevertheless, several caveats limit current TF-DNA structure predictions. Performance varied substantially across TFs, with ETS factors and SF-1 showing strong agreement with SNP-SELEX preferences, while PAX5 and MEIS2 were poorly captured. This heterogeneity likely arises from multiple sources, including biases towards AF3 training templates, the use of JASPAR motifs to guide TF binding sites, and the intrinsic complexity of TFs that bind cooperatively or via multiple domains. Currently, ipTM values can be an unreliable measure of complex model quality (47, 48), as models with low overall confidence, derived for experimentally confirmed interactors, still yield meaningful oligomer preference predictions. The limitations of structural model prediction for complexes containing nucleic acids are well known (36–38) and may be alleviated through expanded crystallographic datasets of TF-DNA interactions, better distillation datasets for bound DNA motifs, as well as confidence metrics tailored to TF-DNA complex predictions. Moreover, the SNP-SELEX context, while ideal to assess physics-based frameworks, oversimplifies transcription factor and DNA interactions. *In vitro* exploration of 40bp oligos does not account for chromatin state, DNA methylation, or cooperative interactions that influence TF binding and non-coding variant effects *in vivo*(29). These limitations highlight that current AF3-based predictions should be interpreted with caution and considered as *post hoc* computational validation rather than guidance for experimental design.

Our study design is also limited by computational resource constraints. While structures representing a single protein conformation have proven to be effective in assessing missense variant effects (32, 58), and in this work we demonstrate a single AF3 prediction contains sufficient information to largely recapitulate SNP-SELEX results for non-coding variants, TF-DNA binding differs fundamentally from protein–protein interactions. Unlike protein–protein interfaces, which are often high-affinity and structurally constrained, TF–DNA interactions are generally weaker, transient, and highly context-dependent (59–61). While some TFs exhibit strong binding to consensus motifs, many interactions occur at lower affinity sites and are probabilistic in nature, with occupancy influenced by chromatin state, cooperative partners, and local sequence environment (39, 42). The current approach also does not take into account the contribution from water molecules, which have been shown to be important for mediating TF-DNA interactions and their specificity (62–65). As such, our single-seed predictions do not reveal whether AF3 can capture the probabilistic binding site landscape or the prediction variability for a given input. Performing multi-seed AF3 runs could provide an auxiliary approach to assessing its confidence by assessing consistency in predicted binding sites, fluctuations in ipTM and how it deals with multiple low-affinity sites or palindromic sequences, which may underly poor performance on PAX5 and MEIS2.

Our findings represent a proof-of-concept for structure-guided assessment of non-coding SNVs that alter TF binding. As AF3 and successor models improve in complex prediction, and address the limitations identified in this work, scaling structural analyses to genome-wide variant datasets may become feasible and practical. Hybrid frameworks that integrate structural physics-based scores with sequence-based predictors could combine accuracy with mechanistic interpretability, offering particular value for rare or clinically relevant variants where training data are scarce, or for TF assemblies where multimeric structural context is critical. We hope this work provides valuable insight to researchers working across diverse fields, from structural model prediction to clinical evaluation of variant effects.

## Methods

### Data prioritization and structural complex model prediction

Original batch SNP-SELEX data was retrieved from GVATdb (23). The 40bp oligos with a variant at the 21st position on the forward strand were extracted using the ‘BSgenome’ R package, based on the coordinates provided in the hg19 reference genome. Due to computational resource constraints, transcription factors (TFs) explored in this work had to be prioritized. We first ranked viable targets based on the number of significant preferentially-bound SNPs (pbSNPs) identified in the SNP-SELEX experiment for each TF. To speed up AF3 predictions we chose to only derive TF-DNA complexes containing a single protein chain. In turn, as the SNP-SELEX experiment allows formation of protein complexes, to ensure the highest possible concordance between the assay values and predictive metrics, we attempted to identify TFs more likely to only bind DNA as monomers through a survey of existing literature (39) and UniProt (66) entries.Using structural TF family annotation on DNA binding preferences (predominantly monomeric *vs* multimeric) from Jolma *et al*., 2013 (39) we narrowed down the viable target list. Finally, we considered the per-TF deltaSVM (23, 28) model performance demonstrated previously, considering examples of TFs with high confidence models and those that did not pass the validation due to low performance. In the end, we selected: 3 ETS family TFs (SPIB, ETV4, ELK3), PAX5 from the PAX family and SF-1 (NR5A1) from the nuclear receptor family due to their described binding of DNA as monomers (40, 67). MEIS2 was also selected for exploration due to its association with the homeobox family. Only SPIB, ELK3 and SF-1 had high-confidence deltaSVM models (AUPRC > 0.75).

Full-length TF sequences were used for all targets except PAX5, for which we used the DBD sequence, retrieved from the SNP-SELEX supplementary files, as the full-length PAX5 assays did not pass the quality control in the original experiment. A local version of AlphaFold 3 was run on a single RTX 6000 Ada (CUDA version 12.6) to derive 75,134 complex models (monomeric TF bound to a DNA duplex) encompassing both reference and alternative allele oligos from all SNP pairs for the six TFs. A seed of ‘1’ was used in the predictions.

For the clinical variant examples, models were predicted using the AlphaFold Server with a random seed. In all cases except the USF2A-USF1 dimer, the oligomeric sequences were derived by retrieving a 41bp window (20 nucleotides in either direction) around the SNV, to more closely match our tested SNP-SELEX set-up. USF2A-USF1 complexes were modelled with 81bp sequences (variant positions with 40bp flanking sequences), as used in Plaisancié *et al*., 2025 (51). Reference genomes used to retrieve the sequences corresponded to those used in the publications (50–54).

### Derivation of structural preference metrics

CIF files for the top predicted complex models were converted to PDBs using BeEM (68) for downstream analyses. FoldX (35) was then used on reference and alternative allele structures to evaluate the energetic impact of DNA mutations. Before proceeding, all complex structures were run through the ‘RepairPDB’ procedure.

In the first instance, the impact of the DNA variants was assessed using ‘BuildModel’, by generating and scoring both the reference-to-alternative (forward) and the alternative-to-reference (reverse) mutations, mutating the respective starting model structures. To carry out DNA mutations in ‘BuildModel’ nucleotides must be specified in lower-case (‘a’,’c’,’t’ or ‘g’). For example, given a structure where B and C chains are a DNA duplex, to mutate the 21st nucleotide on the B chain (which will also mutate the paired nucleotide on C chain, in parallel) the ‘individual_list’ file should read ‘tB21c;’. Here ‘t’ is the reference nucleotide at position 21 on DNA chain B and ‘c’ is the nucleotide being mutated to. FoldX identifies the paired nucleotide on chain C and mutates it correspondingly. The scores were then used to calculate the “symmetric” ΔΔG (ΔΔG_sym_, Eq. 1).

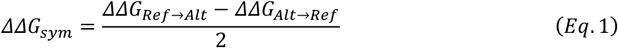

In the second instance, we used the ‘AnalyseComplex’ procedure on the reference and alternative model structures to derive interface strength scores and derived a difference as the ΔInterface metric (Eq. 2).

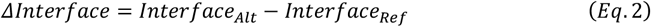

Finally, we also extracted the ipTM scores for each model structure from the AF3 json summary files and derived the difference score for all paired alleles (Eq. 3).

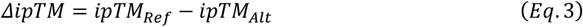

ΔΔG_max(ipTM)_ was derived simply as the ΔΔG score of the model with the higher ipTM in an allele pair, with the reference model score representing a reference-to-alternative mutation, and the alternative score being derived on the alternative model with an alternative-to-reference ‘BuildModel’ mutation (Eq. 4).

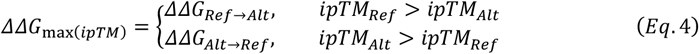

For the TF binding site difference analyses we used FreeSASA(69) to derive 80 element long vectors of per-nucleotide solvent accessible surface area (SASA), encompassing both of the DNA chains in a model. FreeSASA was run with “--hetatm --shrake-rupley --radii=naccess -n 200 --format=seq --n- threads=4” flags. Spearman’s correlation was calculated using base R between the reference and alternative SASA vectors for each SNP pair to identify differences in TF binding position (‘DNA SASA cor.’).

DeltaSVM scores were derived locally using the ‘run.sh’ script provided at https://github.com/ren-lab/deltaSVM. The 94 high-quality models provided for the script only encompassed SPIB, SF-1 and ELK3.

ipSAE (47) was calculated using the script provided at https://github.com/dunbracklab/IPSAE. ‘Mean ipSAE’ represents the average of maximal ipSAE values between the three chains (monomeric TF protein and the DNA duplex). ‘Max ipSAE’ represents the single largest ipSAE observed between interactions of protein and DNA chains only. Values were calculated using 10Å and 15Å distance cutoff values.

## Statistical analysis

The dataset was annotated in line with the system used in the SNP-SELEX paper, where pbSNP pairs were designated as those where the preferential binding score (PBS) p-values were below 0.01 and non-pbSNPs with no differential binding between the pair had p-val > 0.5. pbSNPs with positive PBS values were denoted as showing reference allele preference, while those with negative PBS showed alternative allele preference. The structural metric directions were adjusted to match this convention in the analyses.

Two group comparisons involved a two-sided Mann-Whitney *U* test. In the case of more groups pairwise statistical comparisons were carried out using a Dunn’s test implementation in the R ‘ggstatsplot’ (70) package. The *p*-values for comparisons involving more than two groups were adjusted through Holm’s multiple comparison correction.

We explored the DNA SASA cor. distributions for the pb and non-pb SNPs using the ‘gghistostats’ function from ‘ggstatsplot’. Based on two separate peaks within the distributions, we designated 0.8 as the threshold to separate pairs as those with a ‘matching’ site and those with shifted TF binding sites. We further explored the per-TF density distributions between pairs with pbSNPs and non-pbSNP pairs, statistically assessing differences in distributions using a two-sided Mann-Whitney *U* test. Reported *p*-values were adjusted for multiple testing across TFs using the Benjamini–Hochberg (BH) procedure to control the false discovery rate, using the ‘rstatix’ package.

The overall agreement between the PBS scores and our metrics was assessed through Spearman’s correlation using base R. The full variant pair dataset was used, including the intermediate PBS p-val pairs that were unclassified in the SNP-SELEX experiment.

Two types of classification performance benchmarks were carried out using the ‘PRROC’ (71, 72) package, calculating receiver operating characteristic (ROC) and precision-recall (PR) curves, with the area under the curves (AUC) serving as the performance metric. In the first instance, the metrics were evaluated for capacity to distinguish pbSNPs from non-pbSNPs, with the former as the positive class. We also evaluated how well the metrics can distinguish the different directions of preference (reference allele preference vs alternative allele preference) within only the pbSNPs. In this case PR analysis was performed twice, with each class as the positive. PR-AUC confidence intervals were derived through 1,000 bootstraps, using the ‘boot’ package. Additionally, the performance in distinguishing preference direction was assessed by Matthews correlation coefficient (weighted by |PBS|) through the ‘mcc’ function from the ‘yardstick’ package.

Tests were visualized using ‘ggstatsplot’ and ‘ggplot2’.

## Data Availability

The data generated in this study have been deposited in the OSF database at https://doi.org/10.17605/OSF.IO/UZKG5.

## Acknowledgement

This project was supported by funding from the European Research Council (ERC) under the European Union’s Horizon 2020 research and innovation programme (grant agreement No. 101001169) and by funding from the Medical Research Council (MRC) Human Genetics Unit core grant (MC_UU_00035/9).

## References

1. Lambert, S.A., Jolma, A., Campitelli, L.F., Das, P.K., Yin, Y., Albu, M., Chen, X., Taipale, J., Hughes, T.R. and Weirauch, M.T. (2018) The Human Transcription Factors. Cell, 172, 650–665.

2. French, J.D. and Edwards, S.L. (2020) The Role of Noncoding Variants in Heritable Disease. Trends in Genetics, 36, 880–891.

3. Zhang, F. and Lupski, J.R. (2015) Non-coding genetic variants in human disease. Human Molecular Genetics, 24, R102–R110.

4. Lee, T.I. and Young, R.A. (2013) Transcriptional Regulation and Its Misregulation in Disease. Cell, 152, 1237–1251.

5. Iñiguez-Muñoz, S., Llinàs-Arias, P., Ensenyat-Mendez, M., Bedoya-López, A.F., Orozco, J.I.J., Cortés, J., Roy, A., Forsberg-Nilsson, K., DiNome, M.L. and Marzese, D.M. (2024) Hidden secrets of the cancer genome: unlocking the impact of non-coding mutations in gene regulatory elements. Cellular and Molecular Life Sciences, 81, 274.

6. Barrera, L.A., Vedenko, A., Kurland, J.V., Rogers, J.M., Gisselbrecht, S.S., Rossin, E.J., Woodard, J., Mariani, L., Kock, K.H., Inukai, S., et al. (2016) Survey of variation in human transcription factors reveals prevalent DNA binding changes. Science, 351, 1450–1454.

7. Williamson, K.A., Hall, H.N., Owen, L.J., Livesey, B.J., Hanson, I.M., Adams, G.G.W., Bodek, S., Calvas, P., Castle, B., Clarke, M., et al. (2020) Recurrent heterozygous PAX6 missense variants cause severe bilateral microphthalmia via predictable effects on DNA–protein interaction. Genetics in Medicine, 22, 598–609.

8. McDonnell, A.F., Plech, M., Livesey, B.J., Gerasimavicius, L., Owen, L.J., Hall, H.N., FitzPatrick, D.R., Marsh, J.A. and Kudla, G. (2024) Deep mutational scanning quantifies DNA binding and predicts clinical outcomes of PAX6 variants. Molecular Systems Biology, 20, 825–844.

9. Yang, J. and Adli, M. (2019) Mapping and Making Sense of Noncoding Mutations in the Genome. Cancer Research, 79, 4309–4314.

10. Jin, Y., Jiang, J., Wang, R. and Qin, Z.S. (2021) Systematic Evaluation of DNA Sequence Variations on in vivo Transcription Factor Binding Affinity. Frontiers in Genetics, 12.

11. Deplancke, B., Alpern, D. and Gardeux, V. (2016) The Genetics of Transcription Factor DNA Binding Variation. Cell, 166, 538–554.

12. Landrum, M.J., Lee, J.M., Riley, G.R., Jang, W., Rubinstein, W.S., Church, D.M. and Maglott, D.R. (2014) ClinVar: public archive of relationships among sequence variation and human phenotype. Nucleic Acids Research, 42, D980–D985.

13. Spielmann, M. and Mundlos, S. (2016) Looking beyond the genes: the role of non-coding variants in human disease. Human Molecular Genetics, 25, R157–R165.

14. Gloss, B.S. and Dinger, M.E. (2018) Realizing the significance of noncoding functionality in clinical genomics. Experimental & Molecular Medicine, 50, 1–8.

15. Yao, Q., Ferragina, P., Reshef, Y., Lettre, G., Bauer, D.E. and Pinello, L. (2021) Motif-Raptor: a cell type-specific and transcription factor centric approach for post-GWAS prioritization of causal regulators. Bioinformatics, 37, 2103–2111.

16. Wang, Z., Gong, M., Liu, Y., Xiong, S., Wang, M., Zhou, J. and Zhang, Y. (2022) Towards a better understanding of TF-DNA binding prediction from genomic features. Computers in Biology and Medicine, 149, 105993.

17. John, S., Sabo, P.J., Thurman, R.E., Sung, M.-H., Biddie, S.C., Johnson, T.A., Hager, G.L. and Stamatoyannopoulos, J.A. (2011) Chromatin accessibility pre-determines glucocorticoid receptor binding patterns. Nature Genetics, 43, 264–268.

18. Abramov, S., Boytsov, A., Bykova, D., Penzar, D.D., Yevshin, I., Kolmykov, S.K., Fridman, M.V., Favorov, A.V., Vorontsov, I.E., Baulin, E., et al. (2021) Landscape of allele-specific transcription factor binding in the human genome. Nature Communications, 12, 2751.

19. Srivastava, D. and Mahony, S. (2020) Sequence and chromatin determinants of transcription factor binding and the establishment of cell type-specific binding patterns. Biochimica et Biophysica Acta (BBA) - Gene Regulatory Mechanisms, 1863, 194443.

20. Berger, M.F. and Bulyk, M.L. (2009) Universal protein-binding microarrays for the comprehensive characterization of the DNA-binding specificities of transcription factors. Nature Protocols, 4, 393–411.

21. Pantier, R., Chhatbar, K., Alston, G., Lee, H.Y. and Bird, A. (2022) High-throughput sequencing SELEX for the determination of DNA-binding protein specificities in vitro. STAR Protocols, 3, 101490.

22. Riley, T.R., Slattery, M., Abe, N., Rastogi, C., Liu, D., Mann, R.S. and Bussemaker, H.J. (2014) SELEX-seq: A Method for Characterizing the Complete Repertoire of Binding Site Preferences for Transcription Factor Complexes. In Graba, Y., Rezsohazy, R. (eds), Hox Genes: Methods and Protocols. Springer New York, New York, NY, pp. 255–278.

23. Yan, J., Qiu, Y., Ribeiro dos Santos, A.M., Yin, Y., Li, Y.E., Vinckier, N., Nariai, N., Benaglio, P., Raman, A., Li, X., et al. (2021) Systematic analysis of binding of transcription factors to noncoding variants. Nature, 591, 147–151.

24. Rauluseviciute, I., Riudavets-Puig, R., Blanc-Mathieu, R., Castro-Mondragon, J.A., Ferenc, K., Kumar, V., Lemma, R.B., Lucas, J., Chèneby, J., Baranasic, D., et al. (2024) JASPAR 2024: 20th anniversary of the open-access database of transcription factor binding profiles. Nucleic Acids Research, 52, D174–D182.

25. Dunham, I., Kundaje, A., Aldred, S.F., Collins, P.J., Davis, C.A., Doyle, F., Epstein, C.B., Frietze, S., Harrow, J., Kaul, R., et al. (2012) An integrated encyclopedia of DNA elements in the human genome. Nature, 489, 57–74.

26. Boytsov, A., Abramov, S., Aiusheeva, A.Z., Kasianova, A.M., Baulin, E., Kuznetsov, I.A., Aulchenko, Y.S., Kolmykov, S., Yevshin, I., Kolpakov, F., et al. (2022) ANANASTRA: annotation and enrichment analysis of allele-specific transcription factor binding at SNPs. Nucleic Acids Research, 50, W51–W56.

27. Stormo, G.D. (2000) DNA binding sites: representation and discovery. Bioinformatics, 16, 16–23.

28. Lee, D., Gorkin, D.U., Baker, M., Strober, B.J., Asoni, A.L., McCallion, A.S. and Beer, M.A. (2015) A method to predict the impact of regulatory variants from DNA sequence. Nature Genetics, 47, 955–961.

29. Han, D., Li, Y., Wang, L., Liang, X., Miao, Y., Li, W., Wang, S. and Wang, Z. (2024) Comparative analysis of models in predicting the effects of SNPs on TF-DNA binding using large-scale in vitro and in vivo data. Briefings in Bioinformatics, 25, bbae110.

30. Zhou, J. and Troyanskaya, O.G. (2015) Predicting effects of noncoding variants with deep learning– based sequence model. Nature Methods, 12, 931–934.

31. Tognon, M., Kumbara, A., Betti, A., Ruggeri, L. and Giugno, R. (2025) Benchmarking transcription factor binding site prediction models: a comparative analysis on synthetic and biological data. Briefings in Bioinformatics, 26, bbaf363.

32. Gerasimavicius, L., Livesey, B.J. and Marsh, J.A. (2023) Correspondence between functional scores from deep mutational scans and predicted effects on protein stability. Protein Science, 32, e4688.

33. Delgado, J., Reche, R., Cianferoni, D., Orlando, G., van der Kant, R., Rousseau, F., Schymkowitz, J. and Serrano, L. (2025) FoldX Force Field revisited, an improved version. Bioinformatics, 10.1093/bioinformatics/btaf064.

34. Blanco, J.D., Radusky, L., Climente-González, H. and Serrano, L. (2018) FoldX accurate structural protein–DNA binding prediction using PADA1 (Protein Assisted DNA Assembly 1). Nucleic Acids Research, 46, 3852–3863.

35. Delgado, J., Radusky, L.G., Cianferoni, D. and Serrano, L. (2019) FoldX 5.0: Working with RNA, small molecules and a new graphical interface. Bioinformatics, 35, 4168–4169.

36. Abramson, J., Adler, J., Dunger, J., Evans, R., Green, T., Pritzel, A., Ronneberger, O., Willmore, L., Ballard, A.J., Bambrick, J., et al. (2024) Accurate structure prediction of biomolecular interactions with AlphaFold 3. Nature, 630, 493–500.

37. Krishna, R., Wang, J., Ahern, W., Sturmfels, P., Venkatesh, P., Kalvet, I., Lee, G.R., Morey-Burrows, F.S., Anishchenko, I., Humphreys, I.R., et al. Generalized biomolecular modeling and design with RoseTTAFold All-Atom. Science, 384, eadl2528.

38. Baek, M., McHugh, R., Anishchenko, I., Jiang, H., Baker, D. and DiMaio, F. (2024) Accurate prediction of protein–nucleic acid complexes using RoseTTAFoldNA. Nature Methods, 21, 117–121.

39. Jolma, A., Yan, J., Whitington, T., Toivonen, J., Nitta, K.R., Rastas, P., Morgunova, E., Enge, M., Taipale, M., Wei, G., et al. (2013) DNA-Binding Specificities of Human Transcription Factors. Cell, 152, 327–339.

40. Hoivik, E.A., Lewis, A.E., Aumo, L. and Bakke, M. (2010) Molecular aspects of steroidogenic factor 1 (SF-1). Molecular and Cellular Endocrinology, 315, 27–39.

41. Dupacova, N., Antosova, B., Paces, J. and Kozmik, Z. (2021) Meis homeobox genes control progenitor competence in the retina. Proceedings of the National Academy of Sciences, 118, e2013136118.

42. Jolma, A., Yin, Y., Nitta, K.R., Dave, K., Popov, A., Taipale, M., Enge, M., Kivioja, T., Morgunova, E. and Taipale, J. (2015) DNA-dependent formation of transcription factor pairs alters their binding specificity. Nature, 527, 384–388.

43. Lu, W., Zhang, J., Rao, J., Zhang, Z. and Zheng, S. (2024) AlphaFold3, a secret sauce for predicting mutational effects on protein-protein interactions. bioRxiv, 10.1101/2024.05.25.595871.

44. Yin, Y., Morgunova, E., Jolma, A., Kaasinen, E., Sahu, B., Khund-Sayeed, S., Das, P.K., Kivioja, T., Dave, K., Zhong, F., et al. (2017) Impact of cytosine methylation on DNA binding specificities of human transcription factors. Science, 356, eaaj2239.

45. Abulude, I.J., Luna, I.C.R., Varela, A.S., Camilli, A., Kadouri, D.E. and Guo, X. (2025) Using AlphaFold-Multimer to study novel protein-protein interactions of predation essential hypothetical proteins in Bdellovibrio. Frontiers in Bioinformatics, Volume 5-2025.

46. Evans, R., O’Neill, M., Pritzel, A., Antropova, N., Senior, A., Green, T., Žídek, A., Bates, R., Blackwell, S., Yim, J., et al. (2022) Protein complex prediction with AlphaFold-Multimer. bioRxiv, 10.1101/2021.10.04.463034.

47. Dunbrack, R.L. (2025) <em>Rēs ipSAE loquunt</em>: What’s wrong with AlphaFold’s <em>ipTM</em> score and how to fix it. bioRxiv, 10.1101/2025.02.10.637595.

48. Varga, J.K., Ovchinnikov, S. and Schueler-Furman, O. (2025) actifpTM: a refined confidence metric of AlphaFold2 predictions involving flexible regions. Bioinformatics, 41, btaf107.

49. Burley, S.K., Bhatt, R., Bhikadiya, C., Bi, C., Biester, A., Biswas, P., Bittrich, S., Blaumann, S., Brown, R., Chao, H., et al. (2025) Updated resources for exploring experimentally-determined PDB structures and Computed Structure Models at the RCSB Protein Data Bank. Nucleic Acids Research, 53, D564–D574.

50. Bhatia, S., Bengani, H., Fish, M., Brown, A., Divizia, M.T., de Marco, R., Damante, G., Grainger, R., van Heyningen, V. and Kleinjan, D.A. (2013) Disruption of Autoregulatory Feedback by a Mutation in a Remote, Ultraconserved PAX6 Enhancer Causes Aniridia. The American Journal of Human Genetics, 93, 1126–1134.

51. Plaisancié, J., Angée, C., Erjavec, E., Raymond-Letron, I., Douet, J.-Y., Goetz, M., Vincent-Delorme, C., Karemaker, I.D., Baltissen, M., Vermeulen, M., et al. (2025) Insights into the FOXE3 Transcriptional Network and Disease Mechanisms from the Investigation of a Regulatory Variant Driving Complex Microphthalmia. bioRxiv, 10.1101/2025.01.13.632782.

52. Peña-Martínez, E.G., Rivera-Madera, A., Pomales-Matos, D.A., Sanabria-Alberto, L., Rosario-Cañuelas, B.M., Rodríguez-Ríos, J.M., Carrasquillo-Dones, E.A. and Rodríguez-Martínez, J.A. (2023) Disease-associated non-coding variants alter NKX2-5 DNA-binding affinity. Biochimica et Biophysica Acta (BBA) - Gene Regulatory Mechanisms, 1866, 194906.

53. Oldridge, D.A., Wood, A.C., Weichert-Leahey, N., Crimmins, I., Sussman, R., Winter, C., McDaniel, L.D., Diamond, M., Hart, L.S., Zhu, S., et al. (2015) Genetic predisposition to neuroblastoma mediated by a LMO1 super-enhancer polymorphism. Nature, 528, 418–421.

54. Huang, Q., Whitington, T., Gao, P., Lindberg, J.F., Yang, Y., Sun, J., Väisänen, M.-R., Szulkin, R., Annala, M., Yan, J., et al. (2014) A prostate cancer susceptibility allele at 6q22 increases RFX6 expression by modulating HOXB13 chromatin binding. Nature Genetics, 46, 126–135.

55. Esmaeeli, R., Bauzá, A. and Perez, A. (2023) Structural predictions of protein–DNA binding: MELD-DNA. Nucleic Acids Research, 51, 1625–1636.

56. Ambrosini, G., Vorontsov, I., Penzar, D., Groux, R., Fornes, O., Nikolaeva, D.D., Ballester, B., Grau, J., Grosse, I., Makeev, V., et al. (2020) Insights gained from a comprehensive all-against-all transcription factor binding motif benchmarking study. Genome Biology, 21, 114.

57. Kathail, P., Bajwa, A. and Ioannidis, N.M. (2024) Leveraging genomic deep learning models for non-coding variant effect prediction. 10.48550/arXiv.2411.11158.

58. Cagiada, M., Johansson, K.E., Valanciute, A., Nielsen, S.V., Hartmann-Petersen, R., Yang, J.J., Fowler, D.M., Stein, A. and Lindorff-Larsen, K. (2021) Understanding the Origins of Loss of Protein Function by Analyzing the Effects of Thousands of Variants on Activity and Abundance. Molecular Biology and Evolution, 38, 3235–3246.

59. Shahein, A., López-Malo, M., Istomin, I., Olson, E.J., Cheng, S. and Maerkl, S.J. (2022) Systematic analysis of low-affinity transcription factor binding site clusters in vitro and in vivo establishes their functional relevance. Nature Communications, 13, 5273.

60. Marcovitz, A. and Levy, Y. (2011) Frustration in protein–DNA binding influences conformational switching and target search kinetics. Proceedings of the National Academy of Sciences, 108, 17957–17962.

61. Kribelbauer, J.F., Rastogi, C., Bussemaker, H.J. and Mann, R.S. (2019) Low-Affinity Binding Sites and the Transcription Factor Specificity Paradox in Eukaryotes. Annual Review of Cell and Developmental Biology, 35, 357–379.

62. Morgunova, E., Yin, Y., Das, P.K., Jolma, A., Zhu, F., Popov, A., Xu, Y., Nilsson, L. and Taipale, J. (2018) Two distinct DNA sequences recognized by transcription factors represent enthalpy and entropy optima. eLife, 7, e32963.

63. Morgunova, E., Nagy, G., Yin, Y., Zhu, F., Nayak, S.P., Xiao, T., Sokolov, I., Popov, A., Laughton, C., Grubmuller, H., et al. (2025) Interfacial water confers transcription factors with dinucleotide specificity. Nature Structural & Molecular Biology, 32, 650–661.

64. Li, S. and Bradley, P. (2013) Probing the role of interfacial waters in protein–DNA recognition using a hybrid implicit/explicit solvation model. Proteins: Structure, Function, and Bioinformatics, 81, 1318–1329.

65. Janin, J. (1999) Wet and dry interfaces: the role of solvent in protein–protein and protein–DNA recognition. Structure, 7, R277–R279.

66. The UniProt Consortium (2025) UniProt: the Universal Protein Knowledgebase in 2025. Nucleic Acids Research, 53, D609–D617.

67. Xu, W., Rould, M.A., Jun, S., Desplan, C. and Pabo, C.O. (1995) Crystal structure of a paired domain-DNA complex at 2.5 å resolution reveals structural basis for pax developmental mutations. Cell, 80, 639–650.

68. Zhang, C. (2023) BeEM: fast and faithful conversion of mmCIF format structure files to PDB format. BMC Bioinformatics, 24, 260.

69. Mitternacht, S. (2016) FreeSASA: An open source C library for solvent accessible surface area calculations. 10.12688/f1000research.7931.1.

70. Patil, I. (2021) Visualizations with statistical details: The ‘ggstatsplot’ approach. Journal of Open Source Software, 6, 3167.

71. Grau, J., Grosse, I. and Keilwagen, J. (2015) PRROC: computing and visualizing precision-recall and receiver operating characteristic curves in R. Bioinformatics, 31, 2595–2597.

72. Keilwagen, J., Grosse, I. and Grau, J. (2014) Area under Precision-Recall Curves for Weighted and Unweighted Data. PLOS ONE, 9, e92209.

