## Supplementary for "A structure-guided approach to non-coding variant evaluation for transcription factor binding using AlphaFold 3"

### Supplementary Table 1. Clinical non-coding variant example model sequences.

| TF | SNV | Protein sequence | Forwards strand reference DNA sequence | Forwards strand SNV DNA sequence |
|---|---|---|---|---|
| PAX6 DBD | chr11:31685945 G>T | MQNSHSGVNQLGGVFVNGRPLPDSTRQKIVELAHSGARPCDISRILQVSNGCVSKILGRYYETGSIRPRAIGGSKPRVATPEVVSKIAQYKRECPSIFAWEIRDRLLSEGVCTNDNIPSVSSINRVLRNLASEKQQMGAD | CAAGTGTCAATGCCTGAAGTGATACGCTCTGACAAATCTAA | CAAGTGTCAATGCCTGAAGTTATACGCTCTGACAAATCTAA |
| USF2A | rs745674596 G>A | MDMLDPGLDPAASATAAAAASHDKGPEAEEGVELQEGGDGPGAEEQTAVAITSVQQAAFGDHNIQYQFRTETNGGQVTYRVVQVTDGQLDGQGDTAGAVSVVSTAAFAGGQQAVTQVGVDGAAQRPGPAAASVPPGPAAPFPLAVIQNPFSNGGSPAAEAVSGEARFAYFPASSVGDTTAVSVQTTDQSLQAGGQFYVMMTPQDVLQTGTQRTIAPRTHPYSPKIDGTRTPRDERRRAQHNEVERRRRDKINNWIVQLSKIIPDCNADNSKTGASKGGILSKACDYIRELRQTNQRMQETFKEAERLQMDNELLRQQIEELKNENALLRAQLQQHNLEMVGEGTRQ | TTCACTGGCCATCTTATGCCCGGATGCTCAGCCGCATCACGTCCGGCCCAGGGCCTGTGAAAAGAGGGCCCAGCCACGCTG | TTCACTGGCCATCTTATGCCCGGATGCTCAGCCGCATCACATCCGGCCCAGGGCCTGTGAAAAGAGGGCCCAGCCACGCTG |
| USF1 | | MKGQQKTAETEEGTVQIQEGAVATGEDPTSVAIASIQSAATFPDPNVKYVFRTENGGQVMYRVIQVSEGQLDGQTEGTGAISGYPATQSMTQAVIQGAFTSDDAVDTEGTAAETHYTYFPSTAVGDGAGGTTSGSTAAVVTTQGSEALLGQATPPGTGQFFVMMSPQEVLQGGSQRSIAPRTHPYSPKSEAPRTTRDEKRRAQHNEVERRRRDKINNWIVQLSKIIPDCSMESTKSGQSKGGILSKACDYIQELRQSNHRLSEELQGLDQLQLDNDVLRQQVEDLKNKNLLLRAQLRHHGLEVVIKNDSN | | |
| NKX2-5 | rs7350789 G>A | MFPSPALTPTPFSVKDILNLEQQQRSLAAAGELSARLEATLAPSSCMLAAFKPEAYAGPEAAAPGLPELRAELGRAPSPAKCASAFPAAPAFYPRAYSDPDPAKDPRAEKKELCALQKAVELEKTEADNAERPRARRRRKPRVLFSQAQVYELERRFKQQRYLSAPERDQLASVLKLTSTQVKIWFQNRRYKCKRQRQDQTLELVGLPPPPPPPARRIAVPVLVRDGKPCLGDSAPYAPAYGVGLNPYGYNAYPAYPGYGGAACSPGYSCTAAYPAGPSPAQPATAAANNNFVNFGVGDLNAVQSPGIPQSNSGVSTLHGIRAW | TACAAGAAAGTCTCTACTTGGGTGGTTCACTGCTTTTATAG | TACAAGAAAGTCTCTACTTGAGTGGTTCACTGCTTTTATAG |
| NKX2-5 | rs61216514 G>A | | TTTGAGCCAATATTCCTTGAGTGCCTCTTTGTGCTAGGCAC | TTTGAGCCAATATTCCTTGAATGCCTCTTTGTGCTAGGCAC |
| GATA3 | rs2168101 C>A | MEVTADQPRWVSHHHPAVLNGQHPDTHHPGLSHSYMDAAQYPLPEEVDVLFNIDGQGNHVPPYYGNSVRATVQRYPPTHHGSQVCRPPLLHGSLPWLDGGKALGSHHTASPWNLSPFSKTSIHHGSPGPLSVYPPASSSSLSGGHASPHLFTFPPTPPKDVSPDPSLSTPGSAGSARQDEKECLKYQVPLPDSMKLESSHSRGSMTALGGASSSTHHPITTYPPYVPEYSSGLFPPSSLLGGSPTGFGCKSRPKARSSTGRECVNCGATSTPLWRRDGTGHYLCNACGLYHKMNGQNRPLIKPKRRLSAARRAGTSCANCQTTTTTLWRRNANGDPVCNACGLYYKLHNINRPLTMKKEGIQTRNRKMSSKSKKCKKVHDSLEDFPKNSSFNPAALSRHMSSLSHISPFSHSSHMLTTPTPMHPPSSLSFGPHHPSSMVTAMG | GGGATCCATTTTAGTGTTATCTAAAAATCAAATCTACATGG | GGGATCCATTTTAGTGTTATATAAAAATCAAATCTACATGG |
| HOXB13 | rs339331 T>C | MEPGNYATLDGAKDIEGLLGAGGGRNLVAHSPLTSHPAAPTLMPAVNYAPLDLPGSAEPPKQCHPCPGVPQGTSPAPVPYGYFGGGYYSCRVSRSSLKPCAQAATLAAYPAETPTAGEEYPSRPTEFAFYPGYPGTYQPMASYLDVSVVQTLGAPGEPRHDSLLPVDSYQSWALAGGWNSQMCCQGEQNPPGPFWKAAFADSSGQHPPDACAFRRGRKKRIPYSKGQLRELEREYAANKFITKDKRRKISAATSLSERQITIWFQNRRVKEKKVLAKVKNSATP | GAACTCTCTCTCCCCAGTTTTATGAGGTTTATCTTTAGTGA | GAACTCTCTCTCCCCAGTTTCATGAGGTTTATCTTTAGTGA |

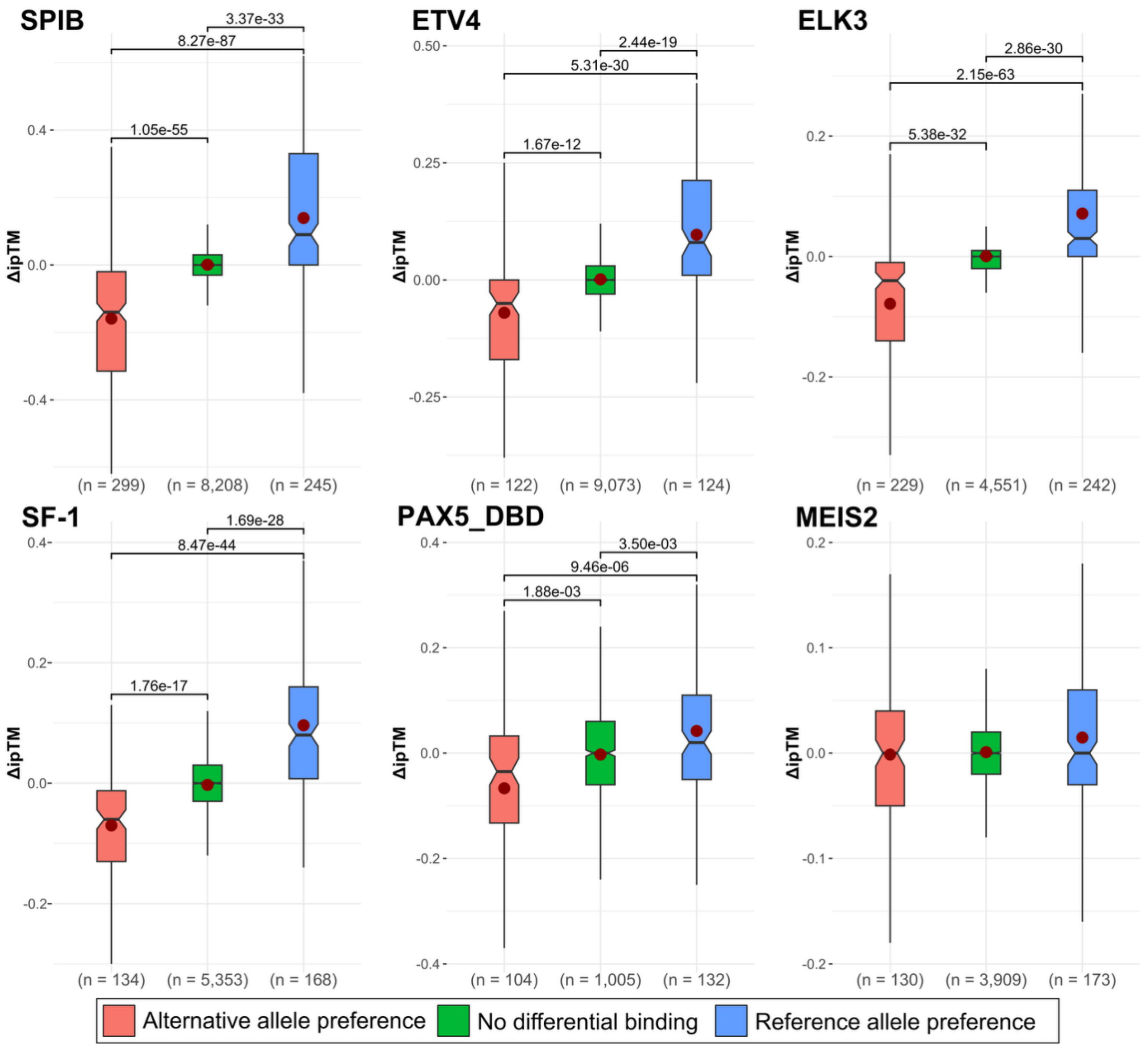


**Supplementary Figure 1. ΔipTM score distributions, in relation to qualitative SNP-SELEX allele preference grouping, reveal TF-specific prediction heterogeneity.** Boxes denote data within 25th and 75th percentiles, and contain median (middle line) and mean (red dot) value notations. Whiskers extend from the box to furthest values within 1.5x the inter-quartile range. Significance values are derived using two-sided Holm-corrected Dunn's tests. Sample size (n) indicates the number of allele pairs.

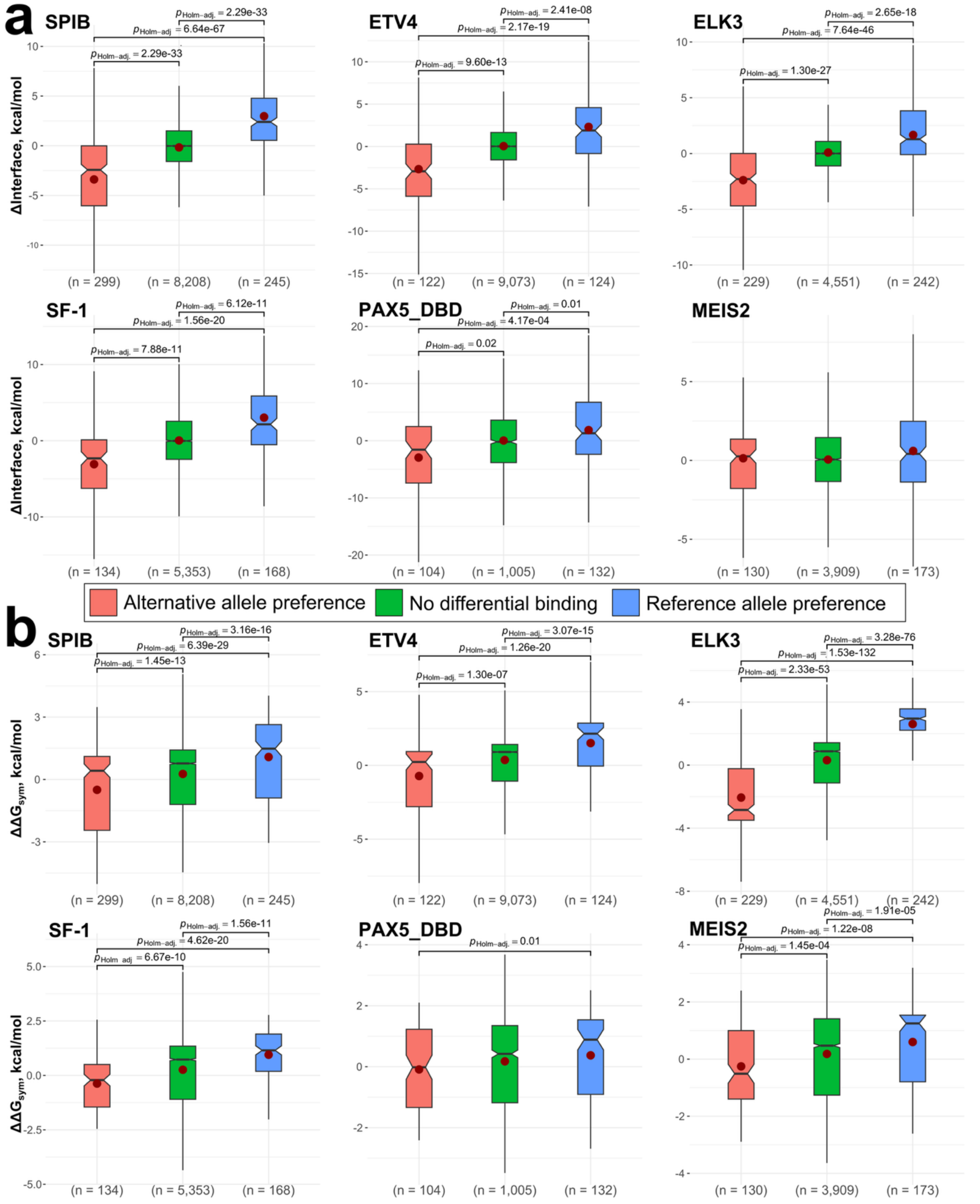


**Supplementary Figure 2. FoldX-derived metrics provide orthogonal information that leads to TF-specific performance improvements. a** – per-TF scores distributions for ΔInterface. **b** – per-TF scores distributions for $\Delta\Delta G_{sym}$. Boxes denote data within 25th and 75th percentiles, and contain median (middle line) and mean (red dot) value notations. Whiskers extend from the box to furthest values within 1.5x the inter-quartile range. Significance values are derived using two-sided Holm-corrected Dunn's tests. Sample size (n) indicates the number of allele pairs.

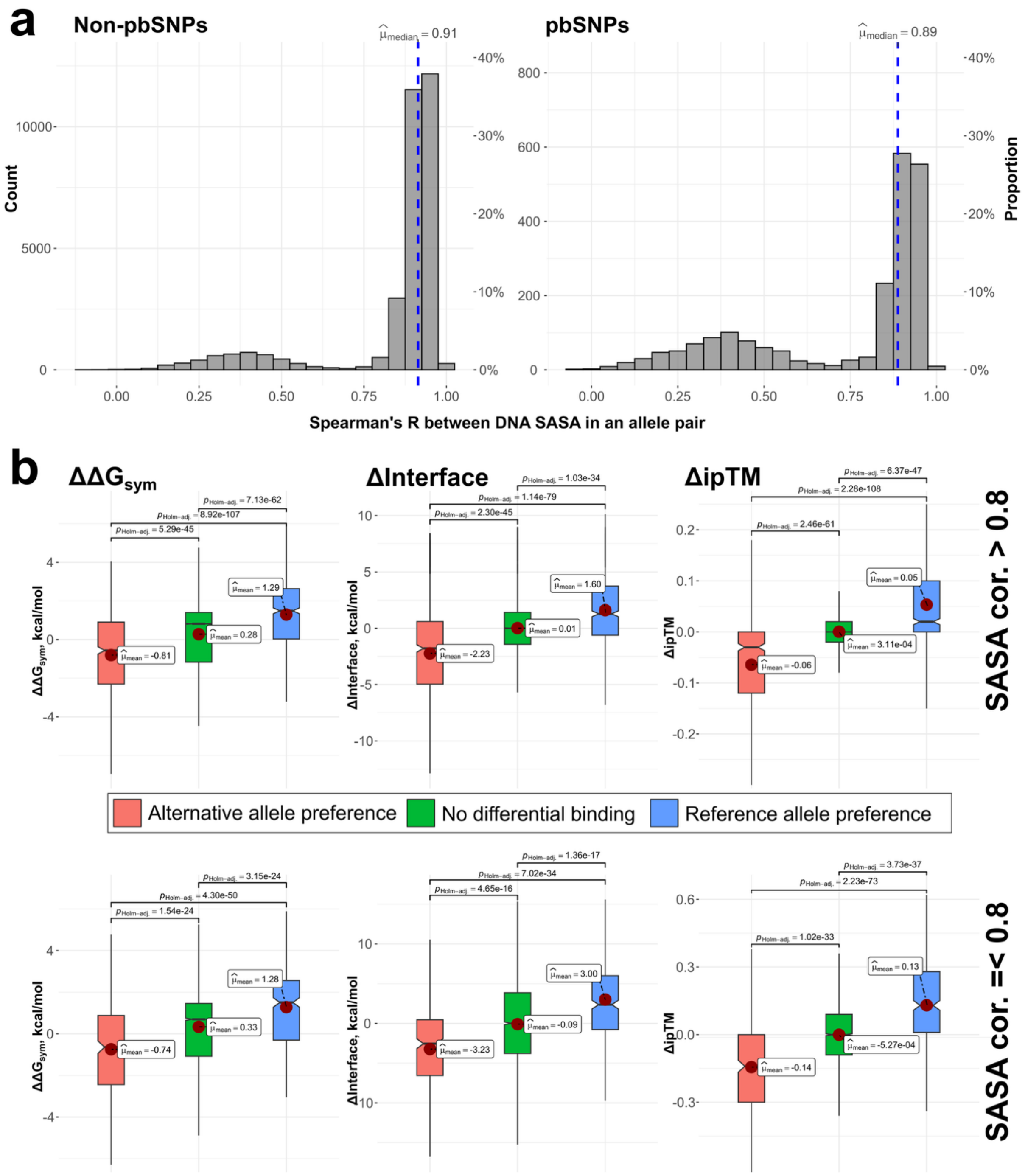


**Supplementary Figure 3. While pairs where one allele is preferentially bound demonstrate more TF binding site shifts, structural metrics still statistically discriminate between SNP-SELEX preference groups in pairs where TFs occupy similar sites.** Analyses represent allele pairs pooled from all six TFs: SPIB, ETV4, ELK3, SF-1, PAX5 DBD and MEIS2. **a** – a histogram of DNA SASA correlation in an allele pair, representative of binding site shifts. **b** – structural score distributions across qualitative SNP-SELEX binding preference groups, based on extent of DNA SASA correlation in an allele pair. Boxes denote data within 25th and 75th percentiles, and contain median (middle line) and mean (red dot) value notations. Whiskers extend from the box to furthest values within 1.5x the inter-quartile range. Significance values are derived using two-sided Holm-corrected Dunn's tests. Sample size (n) indicates the number of allele pairs.

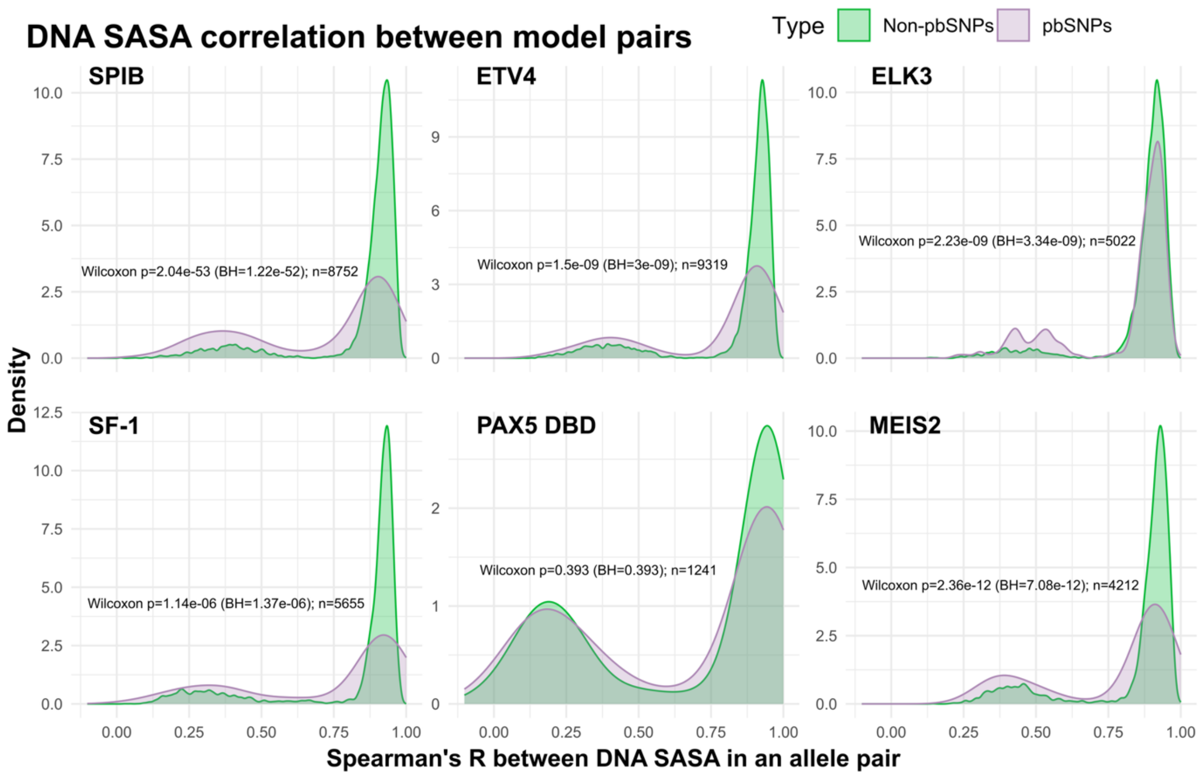


**Supplementary Figure 4. AF3 binding shift behaviour for pbSNP pairs is not universal across TFs.** For each TF, distributions of correlation scores were compared between the two variant groups using the Wilcoxon rank-sum test (also known as the Mann-Whitney *U* test). Reported *p*-values were adjusted for multiple testing across TFs using the Benjamini–Hochberg (BH) procedure[77] to control the false discovery rate.

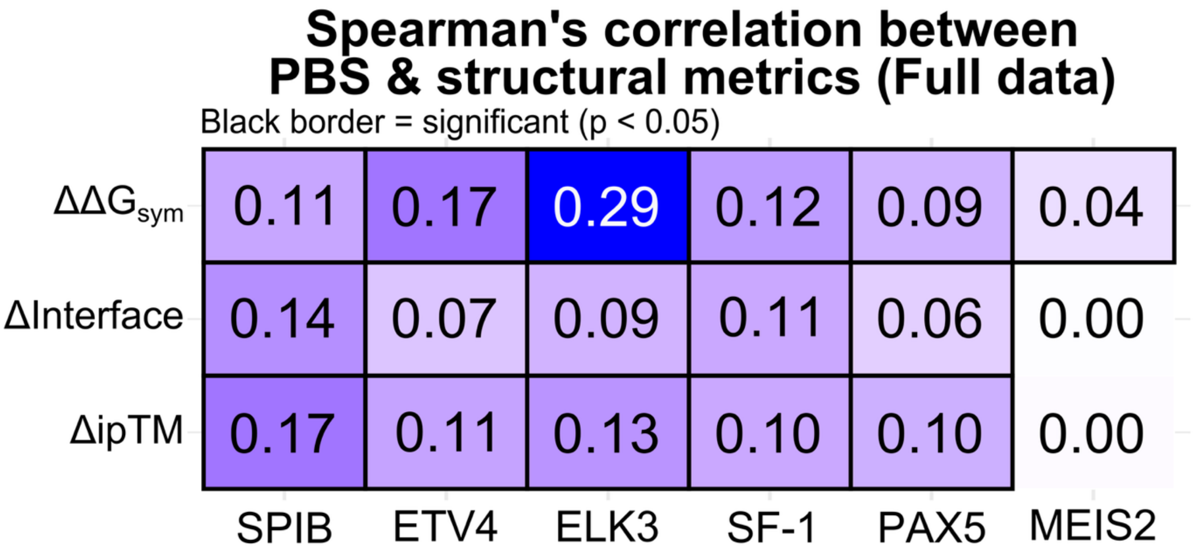


**Supplementary Figure 5. Structural metrics correlate only weakly with the full SNP-SELEX dataset.** Values represent Spearman's correlations between PBS and respective structural metrics, derived per-TF. The full dataset was used, which includes pair intermediate between pbSNPs and non-pbSNPs, which were unclassified in the SNP-SELEX experiment.

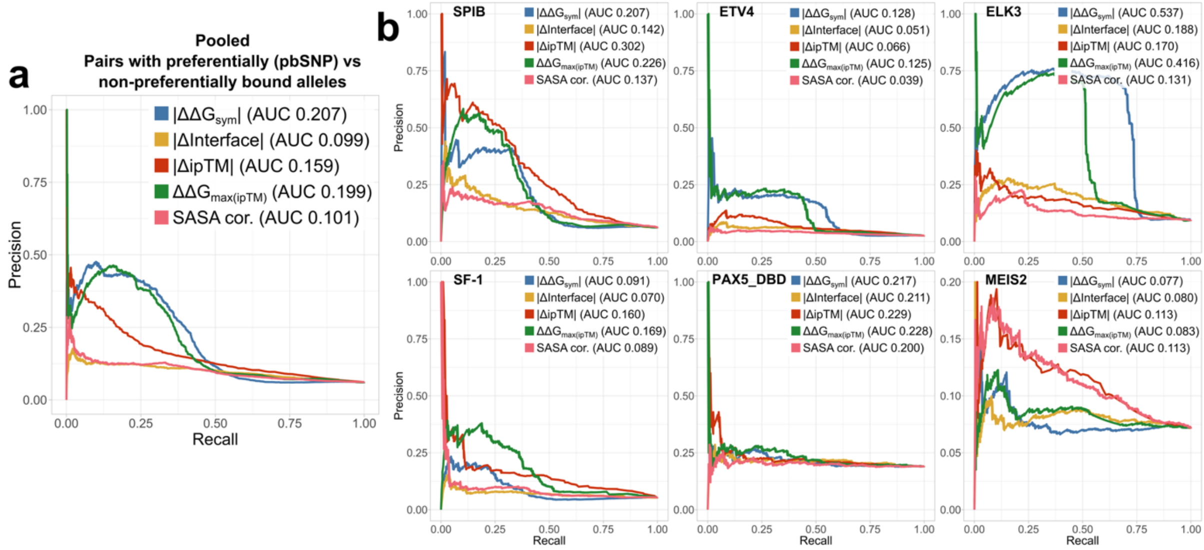


**Supplementary Figure 6. Precision-recall analysis reveals FoldX ΔΔG metrics more accurately identify differentially bound sequence pairs.** Analyses represent precision-recall (PR) curves, with area under the curve (AUC) representing the classification performance. PR analysis explores the identification performance for the positive class, which corresponds to differentially bound (pbSNP) pairs. **a** – Generalized metric performance in distinguishing between pbSNP containing pairs and non-differentially bound sequences (non-pbSNPs). Analysis represents allele pairs pooled from all six TFs: SPIB, ETV4, ELK3, SF-1, PAX5 and MEIS2. **b** – TF-wise classification performance. Qualitative binding preference groups were annotated in line with the SNP-SELEX paper classification. Structural metric directions were adjusted to match the directional convention in the analyses.

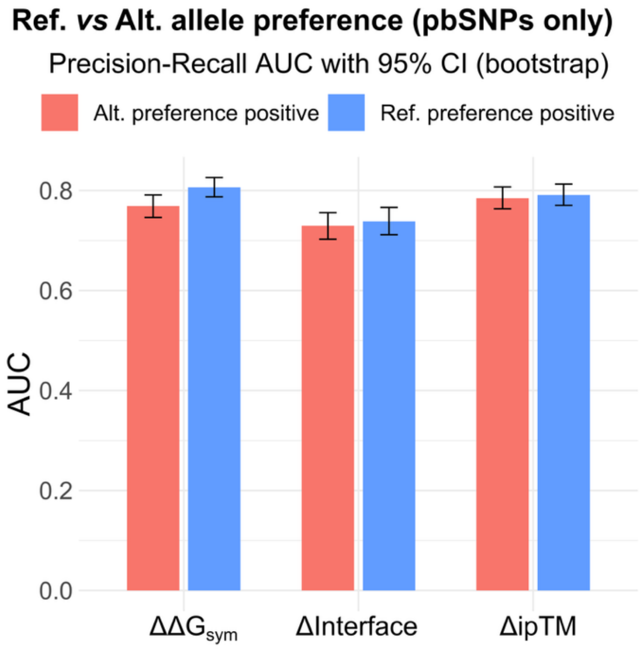


**Supplementary Figure 7. General structural metric classification performance identifies reference and alternative allele binding bias equally well.** Precisions-recall analysis represents allele pairs pooled from all six TFs – SPIB, ETV4, ELK3, SF-1, PAX5 and MEIS2 – with area under the curve (AUC) serving as the classification performance. PR analysis explores the identification performance for the positive class, and due to both directions (*ref.* & *alt.* preference) being informative, it was repeated twice with a different class as the positive.

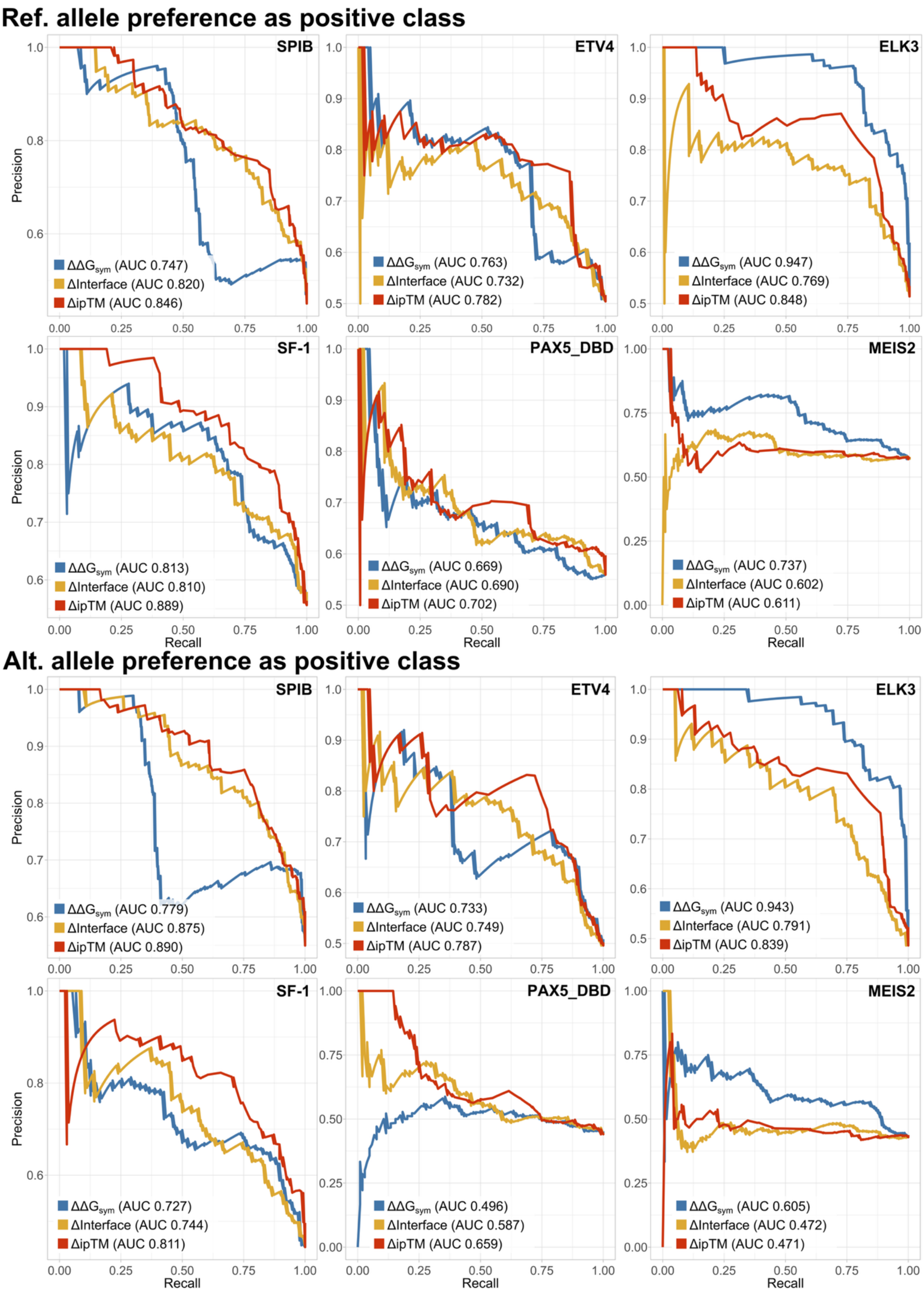


**Supplementary Figure 8. Performance heterogeneity arises when preference direction is explored on a per-TF level.** PR analysis explores the identification performance for the positive class, and due to both directions (*ref.* & *alt.* preference) being informative, it was repeated twice with a different class as the positive.

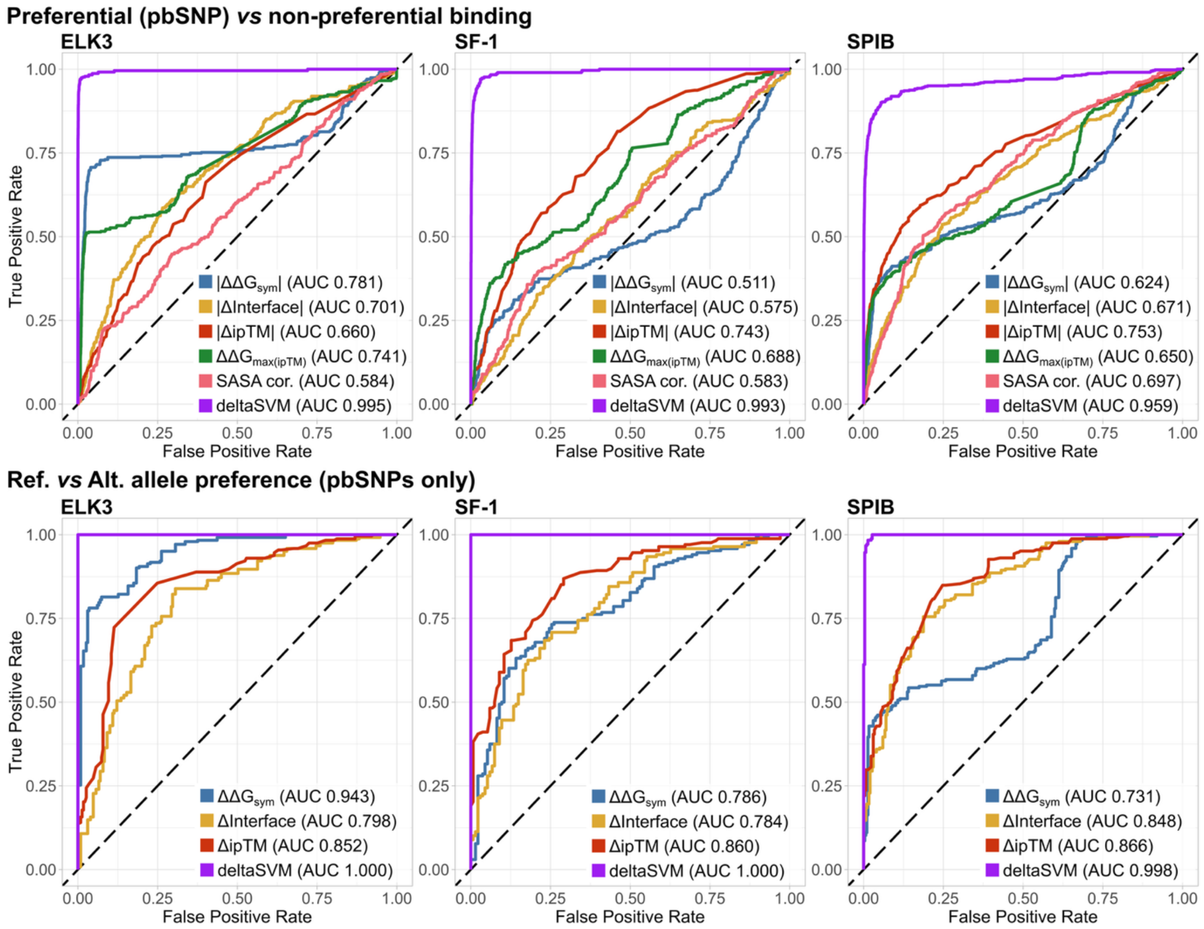


**Supplementary Figure 9. Structural metrics do not achieve state-of-the-art performance presented by deltaSVM models.** Analyses represent receiver operating characteristic (ROC) curves, with area under the curve (AUC) representing the classification performance. ROC analyses were performed for two scenarios: pbSNPs *vs* non-pbSNPS (top); *ref.* preference *vs alt.* preference (pbSNP pairs only). Absolute values were used in the case of pbSNPs *vs* non-pbSNPS for the bi-directional metrics. Metric directions were adjusted to match the directional convention in the analyses.

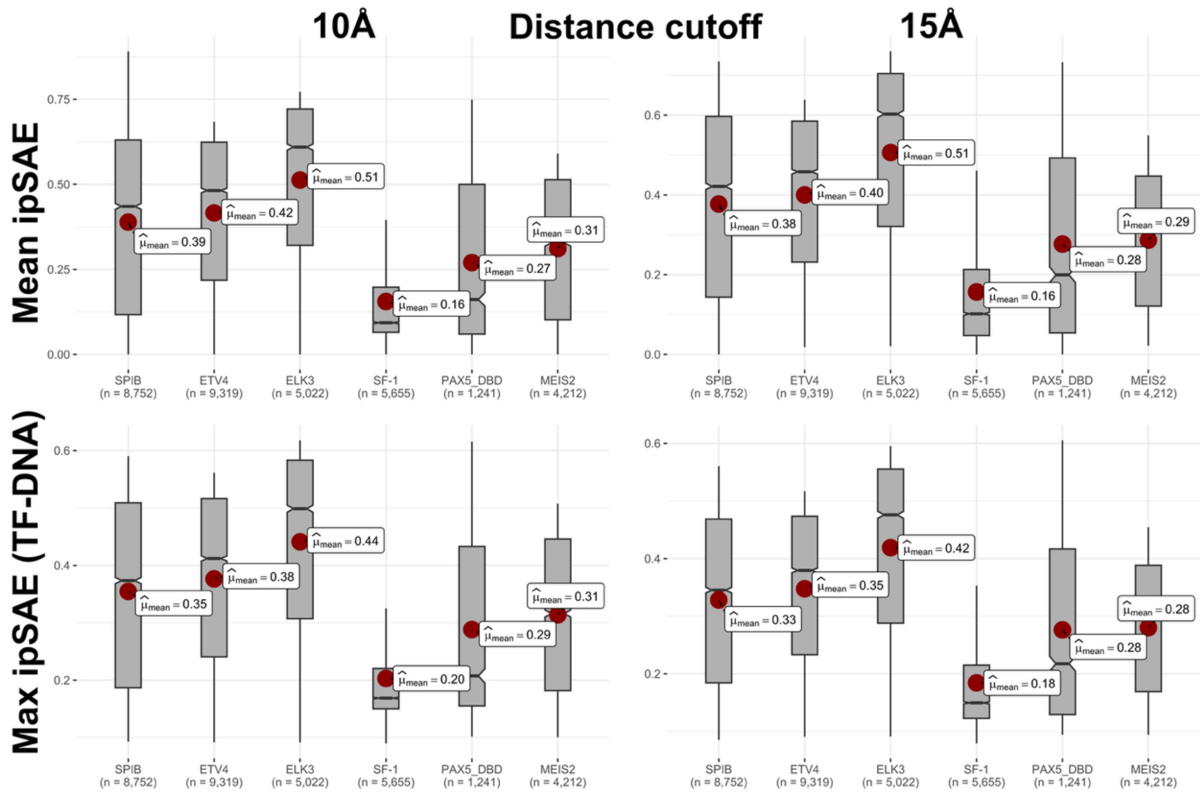


**Supplementary Figure 10. Confidence in TF-DNA model quality does not equalize or improve with an alternative metric.** ‘Mean ipSAE’ represents the average of maximal ipSAE values between the three chains. ‘Max ipSAE’ represents the single largest ipSAE observed between interactions of protein and DNA chains, only. Values were derived using both of the suggested distance cutoffs. Boxes denote data within 25th and 75th percentiles, and contain median (middle line) and mean (red dot) value notations. Whiskers extend from the box to furthest values within 1.5x the inter-quartile range. Sample size (n) indicates the number of allele pairs.

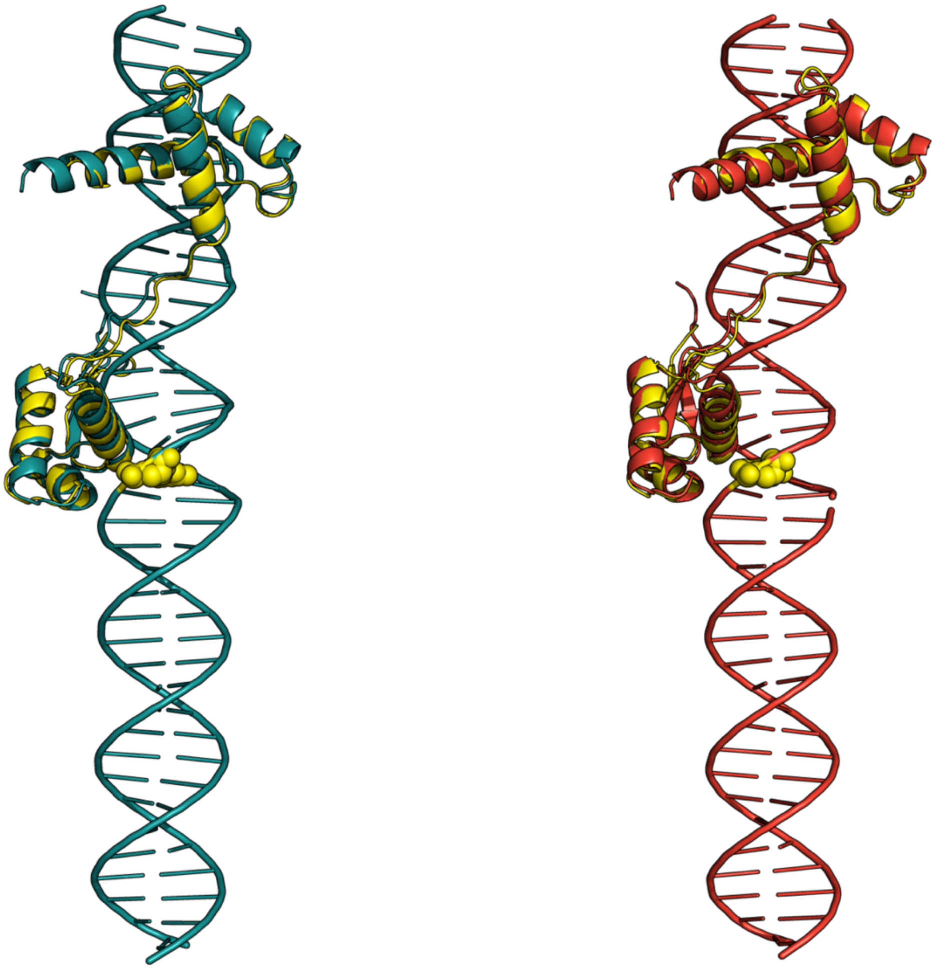

**Supplementary Figure 11. PAX6 paired domain models predicted by AF3 are considerably influenced by PDB templates.** The teal (left) and red (right) models represent the reference and alternative sequence PAX6-DNA models for the chr11:31685945 G>T variant, respectively. Yellow indicates an aligned PAX6 paired domain-DNA complex taken from the PDB (PDB ID 6PAX[85]).
